# High-Affinity Binding of Chemokine Analogs that Display Ligand Bias at the HIV-1 Co-receptor CCR5

**DOI:** 10.1101/575795

**Authors:** Carlos A. Rico, Yamina A. Berchiche, Mizuho Horioka, Jennifer C. Peeler, Emily Lorenzen, He Tian, Manija A. Kazmi, Alexandre Fürstenberg, Hubert Gaertner, Oliver Hartley, Thomas P. Sakmar, Thomas Huber

## Abstract

The chemokine receptor CCR5 is a drug target to prevent transmission of HIV/AIDS. We studied four analogs of the native chemokine RANTES (CCL5) that have anti-HIV potencies of around 25 pM, which is more than four orders-of-magnitude higher than that of RANTES itself. It has been hypothesized that the ultra-high potency of the analogs is due to their ability to bind populations of receptors not accessible to native chemokines. To test this hypothesis, we developed a homogeneous dual-color fluorescence cross-correlation spectroscopy (FCCS) assay for saturation and competition binding experiments. The FCCS assay has the advantage that it does not rely on competition with radioactively labeled native chemokines used in conventional assays. We prepared site-specifically labeled fluorescent analogs using native chemical ligation of synthetic peptides, followed by bioorthogonal fluorescent labeling. We engineered a mammalian cell expression construct to provide fluorescently labeled CCR5, which was purified using a tandem immunoaffinity and size-exclusion chromatography approach to obtain monomeric fluorescent CCR5 in detergent solution. We found subnanomolar binding affinities for the two analogs 5P12-RANTES and 5P14-RANTES, and about twenty-fold reduced affinities for PSC-RANTES and 6P4-RANTES. Using homologous and heterologous competition experiments with unlabeled chemokine analogs, we conclude that the analogs all bind at the same binding site; whereas, the native chemokines (RANTES and MIP1α) fail to displace bound fluorescent analogs even at tens of micromolar concentrations. Our results can be rationalized with *de novo* structural models of the N-terminal tails of the synthetic chemokines that adopt a different binding mode as compared to the parent compound.

## INTRODUCTION

GPCRs are cell surface heptahelical transmembrane receptors that mediate many important physiological processes, and are also involved in the transmission and virulence of several infectious diseases. For example, certain viruses can employ GPCRs during their life cycle to gain cellular entry, enhance dissemination or evade immune detection.(1) The human immunodeficiency virus 1 (HIV-1) utilizes the human CCR5 as a co-receptor to infect immune cells such as dendritic cells and T cells.(2) HIV-1 expresses envelope glycoprotein gp120, which recognizes the cluster of differentiation 4 (CD4) receptor. CD4 engagement causes gp120 to undergo a conformational change that exposes its variable loop 3 (V3), which binds to CCR5. Maraviroc is the only commercial HIV-1 therapeutic agent that targets CCR5 to prevent cellular entry of R5-tropic strains, but case studies have shown the emergence of HIV-1 resistance to maraviroc.(3)

The native chemokine ligands RANTES (Regulated, on Activation, Normal T cell Expressed and Secreted) and MIP-1α (Macrophage Inflammatory Protein 1α) also inhibit HIV-1 entry but have very low potencies.(4, 5) To increase the anti HIV-1 potency chemically modified RANTES analogs were developed(6, 7) that efficiently block viral transmission in macaques.(8) PSC-RANTES, one of these analogs, shows picomolar anti-HIV potency but it is also a strong CCR5 agonist, which may cause undesirable inflammatory effects.(9) Several additional, fully recombinant RANTES analogs were developed using a phage display library by mutating the first nine residues.(9) From this screen, we selected three analogs, 5P12-RANTES (5P12), 5P14-RANTES (5P14), and 6P4-RANTES (6P4) that inhibit HIV-1 with similar potencies as PSC-RANTES (Supplementary Table S1).(9) It is noteworthy that 5P12 is actively developed as a highly effective microbicide candidate in the prevention of human-to-human HIV-1 transmission. Like PSC, 6P4 shows strong agonist activity by calcium flux and internalization assays. In contrast, 5P14 induces receptor internalization but does not activate calcium flux signaling. Last, 5P12 binds to CCR5 but displays no functional activity on CCR5.(9) However, a subsequent study showed that 5P14 can induce Gαi signaling in cyclic adenosine monophosphate (cAMP) inhibition assays in disagreement with earlier conclusions of a lack of G-protein-linked activity.(10) Lorenzen et al. investigated this discrepancy further and showed that 5P12, 5P14, 6P4, PSC, RANTES, and MIP-1α all cause Gαi activation and that PSC and 6P4, and to a lesser extent RANTES and MIP-1α, also induce Gαq activation Based on these data, we proposed a model where the RANTES analogs can bind to both G-protein uncoupled (“naked”) and pre-coupled CCR5 while the native chemokines bind only to the G-protein pre-coupled receptor.(11) Radioligand competition binding assays using ^35^S-gp120 showed that the viral glycoprotein recognizes both the “naked” receptor and the pre-coupled receptor fraction (Colin 2013). Thus, the RANTES analogs efficiently block HIV because they can bind to both the G-protein pre-coupled and the naked CCR5 receptor fractions.

To test the hypothesis that the RANTES analogues can bind to “naked” CCR5 with high affinity, we proposed to develop a strategy where we could measure equilibrium dissociation and competition binding affinities to different receptor fractions. Previously, it was shown that RANTES and MIP-1α require CCR5 G-protein pre-coupling for high affinity binding using a non-hydrolyzable GTP analog that locks the G-protein to the receptor. However, they could not observe the naked receptor fraction using ^125^I-MIP-1α as the tracer. Instead, ^35^S-gp120 was employed since it recognizes both receptor fractions. Yet, the measured chemokine-binding affinities using ^35^S-gp120 were different from the affinities derived using ^125^I-MIP-1α. Because radioligand binding measurements of GPCRs in cell-based systems or crude membrane preparations tend to yield inconsistent results due to their susceptibility to the reaction conditionswe realized that there was an unmet need for a methodology to characterize the ligand binding affinities of the RANTES analogs and the native chemokines with CCR5 in a chemically defined environment.

Single-molecule methods can be used to determine precise receptor-ligand binding parameters of purified components in a chemically defined system. Fluorescence correlation spectroscopy (FCS) is a method that measures the diffusion of a fluorescently-labeled species across an excitation volume. FCS can be used to detect ligand binding of a fluorescently-labeled ligand in the presence of a receptor by observing changes in the ligand’s diffusion time (*τ*_*D*_). If the ligand and receptor are both labeled, then fluorescence cross-correlation spectroscopy (FCCS) can be used to measure the diffusion of the receptor-ligand complex by cross-relating the fluorescence fluctuations from the ligand with the fluorescence fluctuations from the receptor.(12) Absolute concentrations of the ligand, the receptor, and the receptor-ligand complex can also be derived from FCS and FCCS measurements in order to determine equilibrium dissociation constants and inhibition parameters.

To establish a novel FCCS assay to measure ligand-binding parameters of four RANTES analogs with CCR5, we first expressed the CCR5-SNAP construct with a cleavable signal peptide, a SNAP-tag for fluorescent labeling, and N-terminal FLAG and C-terminal 1D4 tags for affinity purification. We then purified monomeric CCR5-SNAP labeled with Alexa-488 (CCR5-SNAP-488) to homogeneity and quantified its concentration by FCS. We prepared the fluorescent chemokines by a modular synthetic scheme that simplifies attachment of different fluorophores to larger peptide ligands. The scheme uses solid-phase peptide synthesis, native chemical fragment ligation, oxime ligation to introduce an azide handle, and strain-promoted azide–alkyne cycloaddition to attach the fluorescent label. Then we used the materials to perform a detailed set of FCCS saturation and competition binding experiments with the CCR5-SNAP-488 and Alexa-647-labeled RANTES analogs. We found that 5P14 and 5P12 bind CCR5 with a very high affinity (sub-nanomolar). In contrast, 6P4 and PSC bind the receptor with more than an order of magnitude lower affinity. Native ligands, such as RANTES, failed to compete at 10 micromolar concentrations with the RANTES analogs binding to CCR5-SNAP-488. We generated homology models of 5P12, 6P4, and RANTES in complex with CCR5 based on the crystal structure of 5P7-RANTES (5P7), another RANTES analog similar to 5P12, bound to CCR5.(13) The homology models revealed that the N-termini of the RANTES analogs bind almost identically to 5P7 in the crystal structure, but differently from RANTES.

## MATERIALS AND METHODS

### Material

Recombinant RANTES and MIP-1α were from PeproTech, Inc. (Rocky Hill, NJ). Coelenterazine 400A for BRET^2^ experiments was from Biotium (Hayward, CA). Forskolin and poly-D-lysine were from Sigma (St. Louis, MO), and the anti-CCR5 mAb (Clone 2D7) directly coupled to phycoerythrin (PE) was from R&D Systems (Minneapolis, MN), anti-CCR5 (Clone T21/8) directly coupled to PE was from eBioscience (San Diego, CA) and anti-Flag PE from Biolegend (San Diego, CA). Dulbecco’s modified Eagle’s medium Glutamax (DMEM-Q), 1% penicillin-streptomycin and Lipofectamine 2000 were from Life Technologies. BSA fraction V, fatty acid free (FAF) was from EMD Millipore and 96-well white microplates with clear bottom, and 384-well black microplates with clear bottom plates were from Corning. Non-labeled RANTES analogs used in this study were prepared by total chemical synthesis as described previously(9) Soluble CD4 was obtained from the National Institutes of Health AIDS reagent program (Catalog number 7356, Lot number 130168) and monomeric BG 505 gp120 (2G12 purified) was a gift from Dr. John Moore (Weill Cornell, NY). Benzylguanine (BG) Alexa-488 was acquired from New England Biolabs (Ipswich, MA).

### Sequence design and molecular cloning of CCR5-SNAP

We designed a synthetic construct encoding the human CCR5 gene with several functional tags for affinity purification and surface immobilization (TH1006, Supplementary Fig. S1). Upstream the CCR5 sequence, we encoded a 5’-adapter with several restriction sites and a Kozak sequence corresponding to a stretch of untranslated amino acids in-frame with the receptor (Supplementary Table S2). We encoded downstream the CCR5 sequence two OLLAS tags separated from CCR5 by a linker sequence containing two KpnI sites. We also encoded the StrepIII tag and the 1D4 epitope downstream the OLLAS tags as alternate tags for affinity purification and immobilization. After the stop codon, we also encoded a 3’-adapter with two restriction sites. TH1006 was then codon optimized using the codon usage in H. *sapiens* using GeneArt (Invitrogen). We protected the restriction sites in the construct during the codon optimization. TH1006 was inserted into the pcDNA3.1(+) plasmid via double digest with NheI and NotI enzymes. To generate TH1007, which has a smaller linker between CCR5 and the OLLAS tags, from TH1006, we digested TH1006 with KpnI and purified the receptor from the linker fragment. We re-ligated the KpnI digested TH1006 using T4 DNA ligase. TOP10 cells were transformed with the ligated reaction product. Single colonies were screened for the correct re-ligated fragment using DNA sequencing. To introduce the SNAP tag downstream CCR5 and generate TH1008 (Supplementary Fig. S1), we performed a double digest of TH1007 and the pSNAPf plasmid (New England Biolabs) with AgeI and BamHI. BamHI and AgeI flank the SNAPf sequence in the pSNAPf plasmid backbone leading to a 570 bp fragment. The 570 bp fragment encoding the SNAPf sequence was isolated and ligated into the AgeI-BamHI double digested TH1008 backbone using T4 DNA ligase. TOP10 cells were transformed with the ligation reaction and single colonies screened for the correct ligated construct. Western blot analysis of TH1008 expression in HEK293T cells showed that the construct was degraded *in vitro* making it unsuitable for ligand binding measurements by FCCS. To overcome this issue, we devised a strategy to incorporate the 5HT_3A_ serotonin receptor signal peptide and FLAG-tag (SP-FLAG) sequence upstream TH1008. We introduced the signal peptide to induce proper transmembrane insertion of the construct and the FLAG epitope to perform a tandem affinity purification to isolate the full-length receptor from truncations. SP-FLAG was designed with a 5’ and a 3’-adapter that allows us to insert the entire coding sequence into different plasmids. The signal peptide and FLAG sequences were encoded without linker between them. Downstream the FLAG tag, we encoded the first 7 amino acids from CCR5 followed by the MlyI restriction site (Supplementary Table S3). We also introduced the double OLLAS tags and the 1D4 epitope flanked by various restrictions sites so we could swap any functional domain between these epitopes. The construct was also codon optimized for expression in H. *sapiens* using GeneArt (Invitrogen). The construct was inserted into the pUC57 plasmid using double digest with NheI and XmaI and named TH1031 (Supplementary Table S3). To introduce the SP-FLAG sequence into the CCR5 constructs, we digested TH1031 with MlyI and purified the 165 bp fragment using agarose gel electrophoresis. We amplified the SP-FLAG fragment using Invitrogen’s Pfx Platinum polymerase (Grand Island, NY) using the primers listed on Supplementary Table S4. The SP-FLAG fragment was inserted upstream TH1007 using Agilent’s QuikChange Lightning mutagenesis kit (Santa Clara, CA) with slight modifications (Supplementary Fig. S1). Briefly, 25 ng of TH1007 was added to 250 ng of purified SP-FLAG in the presence of dNTPs, QuikChange buffer, and polymerase in 25 µL total volume. The PCR reaction was cycled using the parameters as described by Agilent. XL10-Gold cells were then transformed with the PCR reaction per Agilent’s instructions. Single colonies were isolated and screened for the correct construct using DNA sequencing. The construct derived from TH1007 with the SP-FLAG upstream CCR5 is called TH1040. To introduce SNAPf into TH1040, we performed a double digest of TH1040 and TH1008 using HindIII and KpnI. We resolved the desired DNA fragments using agarose gel electrophoresis and purified them using Millipore’s DNA extraction centrifugal filter units (Billerica, MA). The 1204 bp fragment from TH1040 was inserted into the TH1008 backbone while retaining the sequence in frame. The sequences were ligated using T4 DNA ligase per the manufacturer’s instructions. TOP10 cells were transformed with the ligation reaction and single colonies were isolated and screened for the correct construct using DNA sequencing. The new construct with the SP-FLAG sequence upstream CCR5 in TH1008 is then named TH1030 (Supplementary Fig. S1).

### Cell culture and transfection

HEK293T cells (passage number 5 to 15, ATCC, Manassas, VA) were maintained in DMEM-Q, 1% penicillin-streptomycin, and 10% fetal bovine serum from Atlanta Biologicals (Flowery Branch, GA). Unless otherwise noted, transient transfections including high-throughput in-plate transfections were performed using Lipofectamine 2000 according to manufacturer’s instructions with some modifications as described previously.(14) Total transfected plasmid DNA was kept constant by adding empty vector pcDNA3.1(+) when necessary. The total plasmid DNA in all our experiments was 8 µg in 10 cm dishes, 2 µg in 6-well plates, 100 ng in 96-well plates, 20 ng in 384-well plates.

### Flow cytometry

HEK293T cells were transfected in six-well plates with 0.75 µg CCR5 WT or 2.0 µg CCR5-SNAP or 2 µg of empty vector pcDNA3.1(+). Cells were detached in ice cold PBS. Cells were then distributed in 96 well round bottom plates, spun down and re-suspended in BRET buffer (PBS with 0.5mM MgCl_2_ and 0.1% BSA, Fraction V, fatty acid free) containing, anti-CCR5 mAb (Clone 2D7) or anti-CCR5 PE (Clone T21/8) or anti-FLAG PE for 45 minutes at 4°C. Cells were then washed three times in ice cold PBS. Cell surface expression was quantified by flow cytometry using the Accuri C6 flow cytometer (BD Biosciences).

### Adenylyl cyclase activity

HEK293T cells were co-transfected in a high-throughput in-plate manner with 12 ng RLuc3-EPAC-GFP10, a BRET^2^ cAMP sensor (a gift from Dr. Bouvier, Université de Montréal) and 23 ng CCR5 WT or 60 ng CCR5-SNAP or 88 ng of empty vector pcDNA3.1(+). Cells were then plated into 0.01% poly-D-Lysine coated 96-well, white microplates with clear bottom at a density of 100,000 cells/well. Twenty-four hours post transfection; media was replaced with BRET buffer. Coelenterazine 400A was added at a final concentration of 5 µM followed by a 5 minutes incubation at room temperature. Cells were then stimulated with ligand in the presence or absence of 5 µM of forskolin at room temperature for 5 minutes. Luminescence and fluorescence readings were collected using the Synergy NEO2 plate reader from Biotek (Winooski, USA) and Gen5 software. BRET^2^ readings between Rluc3 and GFP10 were collected by sequential integration of the signals detected in the 365 to 435 nm (Rluc3) and 505 to 525 nm (GFP10) windows. BRET^2^ ratios were calculated as described previously.(15, 16) Dose-response curves were fitted using a 3-parameter logistic in GraphPad Prism. The logistic equation is defined as follows

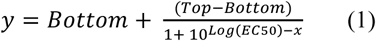

where *x* is the logarithm of agonist concentration, *y* is the response, *bottom* is the bottom plateau, *top* is the top plateau also known as E_max_, and *EC*50 is the effective concentration that yields 50% response.

### Calcium flux assay

For each well of a 384-well plate, 20,000 HEK293T cells in 20 µL DMEM were transfected with 7.5 ng CCR5 WT or 20 ng CCR5-SNAP. Transfected HEK293T cells were plated into 384-well plates (Corning) coated with poly-D-lysine hydrobromide (Sigma-Aldrich, St. Louis, MO) at 20 µL/well. 24 hours post-transfection, 20 µL/well FLIPR calcium 6 dye (Molecular Devices, Sunnyvale, CA) was added to the cells and incubated for 1.5 hours at 37°C with 5% CO_2_. The dye was dissolved in HBSS-H (Hank’s Balanced Salt Solution with 20 mM HEPES, pH 7.4) and supplemented with 0.4% BSA (Fraction V, FAF, Roche). Prior to measurement, the plate was incubated at 37°C for an additional 30 minutes in a pre-warmed FlexStation II 384 Plate Reader (Molecular Devices). Ligands at a 5x final concentration were diluted in HBSS-H supplemented with 0.2% BSA. Fluorescence readings were collected using the FlexStation plate reader with excitation at 485 nm, emission at 535 nm and dichroic mirror at 525 nm. The FlexStation took measurements over a 100 second time course, with 10 µL of ligand added to the cells 20 seconds after the start of measurement. Relative fluorescence units (RFU) are reported as the peak magnitude signal subtracted by the basal signal in each well. Dose-response curves were fitted using the same 3-parameter logistic equation employed to fit the dose-response data from cAMP inhibition experiments.

### Expression, labeling and purification of CCR5-SNAP

Ten 100-mm x 20-mm polystyrene dishes were plated with HEK293T cells at 4.0×10^6^ cells/dish in DMEM + 10% FBS. 24 hours post-plating, 100 µL of Plus Reagent was mixed with 80 µg of CCR5-SNAP in 7.5 mL of DMEM. In a separate vessel, 170 µL of lipofectamine reagent was mixed with 5 mL of DMEM. After 15 minutes, the transfection solutions were mixed and incubated for an additional 15 minutes. Media was removed from HEK293T cells and supplemented with 2.8 mL of DMEM. 1.2 mL of the transfection solution was added to each plate and the cells were incubated for 4 hours before supplementing the media with 4 mL of DMEM + 20% FBS. 24 hours post-transfection, media was removed from the cells and cells were harvested in 2 mL/dish of PBS and 1 mM phenylmethylsulfonyl fluoride (PMSF). Cells were pelleted in a 50 mL vessel at 1,500 rpm using a Beckman GS-6R centrifuge at 4 °C for 5 minutes. The harvesting solution was removed and the cell pellet was solubilized in 5 mL of Buffer L (20 mM HEPES pH 7.4, 0.1 M (NH_4_)_2_SO_4_, 1 mM CaCl_2_, 5 mM MgCl_2_, 10% Glycerol, 0.1% CHS, 1.0% DDM, 1.0% CHAPS) supplemented with a protease inhibitor cocktail for 2 hours at 4 °C. Cell lysates were then centrifuged at 55,000 rpm for 30 minutes, 4 °C, using a TLA 100.3 rotor. The supernatant was added to 600 µL of 50% slurry 1D4 mAb Sepharose 2B resin and incubated overnight at 4 °C. Resin was pelleted in a GS-6R for 5 minutes, 2,000 rpm, 4 °C and then transferred to a Ultrafree-MC-HV Durapore PVDF 0.45 µm centrifugal unit. CCR5-SNAP was labeled in 400 µL of Buffer N (20 mM HEPES pH 7.4, 0.1 M (NH_4_)_2_SO_4_, 1 mM CaCl_2_, 5 mM MgCl_2_, 10% Glycerol, 0.07% CHS, 0.33% DDM, 0.33% CHAPS, 0.018% DOPC, 0.008% DOPS) with 50 µM SNAP-substrate and 1 mM DTT for 30 minutes at room temperature. Resin was then washed 3 x 0.5 mL in Buffer N for 30 minutes each at 4 °C. CCR5-SNAP was eluted from the 1D4 resin by incubating the sample with 1D5 peptide in Buffer N (0.33 mg/mL) twice for 30 minutes on ice and eluting by centrifugation. 1D4 purified CCR5-SNAP was added to 100 µL of FLAG M2 resin and incubated overnight at 4 °C. FLAG resin was transferred to a separate Durapore spin filter and washed 3 times with 0.5 mL of Buffer N for 30 minutes each at 4 °C. CCR5-SNAP was eluted by incubating the resin twice with 100 µL of buffer N and FLAG peptide (200 µg/mL) for 30 minutes on ice. FLAG purified CCR5-SNAP was loaded into a Superdex 200 10/300 GL column previously equilibrated with Buffer N and 0.1 mg/mL BSA (IgG free). CCR5-SNAP was eluted over 1 column volume into 0.5 mL fractions. NIR-immunoblotting and fluorescence correlation spectroscopy were employed to analyze the size exclusion chromatography (SEC) fractions.

### Immunostaining and total internal reflection fluorescence microscopy (TIRF)

HEK293T cells were plated onto 35 mm glass bottom (1.5) Matek plates at 300,000 cells per dish. Cells were transfected with WT CCR5 (0.75 µg), CCR5-SNAP (2.0 µg), or pcDNA3.1(+). (2.0 µg) at a total DNA/dish ratio of 2.0 µg using Lipofectamine 2000 per manufacturer’s instructions. 24 hours post-transfection, media was aspirated from the plates and cells washed with 1 x 2 mL of PBS supplemented with Ca^2+^ and Mg^2+^ (Ca/Mg). Cells were then permeabilized with 1 mL of cold methanol for 5 minutes at −20 °C. Cells were then washed 3×1 mL of cold PBS (Ca/Mg) before blocking overnight in 0.5% BSA in PBS (Ca/Mg) at 4 °C. Blocking solution was removed and 1D4 monoclonal antibody at a dilution of 1:2,000 in 0.5% BSA-PBS (Ca/Mg) was added for one hour at room temperature. Cells were then washed with 3×1 mL of PBS (Ca/Mg). Secondary antibodies conjugated to Alexa-488 were added at a final dilution of 1:500 in 0.5% BSA-PBS (Ca/Mg) for 1 hour at room temperature. Cells were washed again 3×1 mL of PBS (Ca/Mg) and then LiCor mounting media containing DAPI was added to the cells. Cells were visualized on a Nikon TiE inverted TIRF-FLIM microscope using an Apo TIRF 100x oil N2 objective (N.A. 1.49). Images were collected on an Andor NEO sCMOS camera using 405-and 488-nm excitation with a total exposure of 150 ms per image. Images were acquired at room temperature using the following dimension order, XYCZT, which are 2048, 2048, 3, 1, 1 pixels respectively. Filters used were 525/50 and 450/40. Images were processed using ImageJ and Adobe Illustrator.

### SDS-PAGE analysis and NIR-immunoblotting

Samples were mixed with DTT at 150 mM final concentration and NuPAGE loading buffer. Samples were loaded into a NuPAGE 4-12% Bis-Tris gel in MES-SDS buffer. Electrophoresis was conducted at a constant voltage of 115V. The gel was removed from the cassette and rinsed in water before equilibrating in Western Transfer buffer (48 mM Tris, 39 mM glycine, 1.3 mM SDS, 20% MeOH, pH 9.2). 1 piece of Immobilon PVDF membrane-Fl was incubated for 1 minute at room temperature in 100% MeOH. The PVDF membrane and 2 pieces of extra thick blot papers (Bio-Rad) were rinsed in Western transfer buffer. Western transfer was performed in a semi-dry apparatus for 45 minutes with a constant voltage of 18V. After electrophoresis, the membrane was placed in 10 mL of Odyssey blocking buffer (PBS) and incubated for 1 hour at room temperature. The membrane was then placed in 10 mL of blocking buffer with anti-1D4 mouse monoclonal (1:1,000), anti-FLAG rabbit polyclonal (1:1,000) antibodies, and 0.2% Tween-20. The membrane was incubated overnight at 4 °C. Membrane was then washed 5×5 minutes in 1x PBS-T (0.1% Tween-20). Membrane was incubated for 1 hour at room temperature in 10 mL blocking buffer supplemented with 0.2% Tween-20, 0.01% SDS, goat anti-mouse IR 680 RD (1:10,000), and goat anti-rabbit IR 800 CW (1:10,000). Membrane was washed again 5×5 minutes in 1× PBS-T and then 2×5 minutes in 1× PBS buffer. Membranes were visualized using a LICOR Odyssey SA using 100 µm resolution, and intensity level 7 for both 700 and 800 nm excitations. Images were processed using Image Studio Lite Version 4.0 and ImageJ. For the line scan analysis, a rectangle of 45 × 120 pixels was drawn around the desired gel lane and set as First Lane under Analyze, Gels, in ImageJ. The command ‘Plot Lanes’ was then selected with vertical and horizontal scale factors set to 1.0 with uncalibrated optical density. Using the magic wand, an area under the curve was selected and saved as *x* and *y* coordinates for replotting in GraphPad Prism 7.

### Synthesis of fluorescently-labeled chemokines

Fluorescent chemokines were prepared by total chemical synthesis. First, N-terminal and C-terminal fragments corresponding to residue numbers 1-33 and 34-68 respectively were made by polymer-supported organic synthesis using Boc chemistry. The C-terminal fragment was common to all chemokines and consisted of RANTES (34-68). The C-terminal fragment additionally carried the K(S)G sequence, where the serine was coupled to the ε-position of the lysine, as a C-terminal extension. After deprotection, cleavage from the resin, and purification, N-terminal fragments were coupled to the extended C-terminal fragment by native chemical ligation under previously described conditions.(6) Crude products were analyzed by HPLC and ESI-MS before protein refolding and formation of the intramolecular disulfide bridges. The refolded material was then further analyzed by HPLC on a C8 column (shorter retention time; Aeris WIDEPORE 3.6u XB-C8) and ESI-MS (lower mass due to disulfide bridge formation; instrument: Bruker Esquire 3000+ ion trap mass spectrometer).

To introduce a reactive handle for chemical derivatization of the chemokines, the serine residue on the C-terminal extension was selectively oxidized to yield a glyoxylyl residue, whose aldehyde reacts efficiently and specifically with aminooxy compounds. The oxidation was performed with 10 equivalents of NaIO_4_ in 1% NH_4_HCO_3_ buffer at pH 7.2 in the presence of 50 equivalents of methionine.(17) The mixture was left to react in the dark for 10 min before the reaction was quenched by the addition of a 10,000-fold excess of ethylene glycol. After 10 additional minutes, the solution was acidified and the chemokines were isolated on a C18 Sep-Pak column and analyzed by ESI-MS (For PSC, expected: 8134.5; found: 8135.3±0.8; 5P12: expected 8181.7, found 8181.2±0.8; 6P4: expected 8118.5, found 8118.3±0.4; 5P14: expected 8186.0, found 8185.8±0.6; RANTES: expected 8088.3, found 8088.0±0.3). The material was then freeze-dried. In order to prevent oxidation of the N-terminal Serine of RANTES during the procedure, Serine was temporarily protected with a 2-(methylsulfonyl)ethyl oxycarbonyl (Msc) group.(18) The protection was removed by dissolving the compound in a 1:1 water:DMF mixture, cooling it to 0°C, and then adding NaOH at a final concentration of 0.5M. After 30s, acetic acid was added to the solution and the material was isolated again on a C18 Sep-Pak column.

To couple Alexa-647 to the chemokines via strain-promoted azide-alkyne cycloaddition (SpAAC) reaction, a bifunctional linker carrying on one side an aminooxyacetate (AOA) moiety and on the other side an azide was synthesized. First, desalted Boc-protected AOA dicyclohexylamine was converted into its N-hydroxysuccinimidyl (NHS) ester derivative following standard procedures.(19) Briefly, 2.35 mmol of the amino acid were left to react overnight with 2.4 mmol of N-hydroxysuccinimide and 2.4 mmol of N,N’-dicyclohexylcarbodiimide in 10 ml ethyl acetate. The mixture was then filtrated, dried, resuspended in dichloromethane and dried again. 2.23 mmol of Boc-protected AOA NHS ester were recovered. Subsequently, 1 mg of Boc-protected AOA NHS ester in 2 ml of dichloromethane was slowly added to 5 mmol of ethylenediamine (en) and left to react for 3 hours. Boc-AOA-en was then isolated by HPLC on a C8 column (87 mg of product recovered). 75 mg (0.22 mmol) of this compound was further reacted for 24 hours with 72 mg (0.22 mmol) of Boc-3-azidoalanine (Ala(N_3_)) NHS ester (made from Boc-3-azidoalanine and N-hydroxysuccinimide according to the procedure described above) in 4 ml of acetonitrile in the presence of 0.66 mmol N-methylmorpholine. The desired linker with the overall structure Boc-AOA-en-Ala(N_3_)-Boc was isolated by HPLC on a C8 column and its mass confirmed by ESI-MS (expected: 445.8; found: 446.3 (M+H)). Boc-AOA-en-Ala(N_3_)-Boc was then deprotected in trifluoroacetic acid (TFA) for 10 min, dried, analyzed by ESI-MS, and resuspended in water. 20 equivalents of AOA-en-Ala(N_3_) were added to 150 nmol of the different oxidized chemokines at a concentration of 100-200 µM and left to react overnight in 90 mM sodium formate buffer at pH 3.0. The product of the reaction (140 nmol) was isolated on a C18 Sep-Pak column.

DIBO-modified Alexa Fluor 647 (DIBO-AF647, Life Technologies, C10408) was dissolved at 5 mM concentration in DMSO. The chemokines derivatized with the Ala(N_3_) linker were labeled with DIBO-AF647 by incubating a mixture of protein (300-400 µM) with 1.8 reactive dye equivalents and letting the reaction proceed at room temperature overnight. The labeled protein was recovered on an S-2000 column (Phenomenex Biosep-SEC-S-2000) equilibrated in 50% MeOH and its mass confirmed by ESI-MS (PSC: expected 9523.8, found 9524.0±0.5; RANTES: expected 9477.5, found 9476.8±0.5; 6P4: expected 9472.6, found 9472.2±0.2; 5P12: expected 9535.7, found 9535.2±0.3; 5P14: expected 9593.1, found 9591.2±0.3). The absence of a peak in the MS spectrum at the mass of the unlabeled protein further indicated that the labeling reaction was quantitative. Supplementary Figure S2 shows the chemical structure of the C-terminal extension on lysine for all the chemokines synthesized in this study.

### Ligand binding assays

Saturation ligand binding assays were set-up in PCR tubes by serially diluting the ligand in Buffer N supplemented with 0.1 mg/mL BSA (IgG free, detergent free). CCR5-SNAP-A488 was then added in equal volume for a total reaction volume of 20 µL. Samples were equilibrated at room temperature for 4 hours protected from ambient light. 15 µL of each sample were loaded into individual wells of a 384 well plate previously blocked with 1.0 mg/mL BSA (IgG free, detergent free) in water for 15 minutes at room temperature. To prevent sample evaporation, 5 to 10 µL of paraffin oil was applied to the top of each sample. Competition binding assays were set-up in a similar fashion except that the labeled chemokine was kept at constant concentration and the competitor was serially diluted. 5 µL of the labeled chemokine was mixed with 5 µL of non-labeled chemokine and then 10 µL of CCR5-SNAP-A488 was added for a 20 µL total reaction volume. Samples were equilibrated for ≥ 16 hours at room temperature before imaging by FCCS. For competition with the sCD4-gp120 complex, sCD4 and gp120 were incubated for 1 hour at a molar ratio of 10:1 respectively and final complex concentration of 20 µM. Complex was then serially diluted in Buffer N prior to adding labeled 5P12-or 6P4-647 and CCR5-SNAP-488. Samples were then incubated for ≥ 16 hours at room temperature prior to FCCS measurements.

### Fluorescence correlation and cross-correlation spectroscopy measurements

Samples were loaded into # 1.5 glass-bottom 96- or 384-well black plates (SensoPlates, black, 384 well reference number: 788892, 96 well plate reference number: 655892, Greiner bio-one) and mounted in an inverted laser scanning confocal microscope LSM 780 (Zeiss). Alexa-488 was excited using an Argon 488 nm laser at 0.2% or 0.8% laser transmission and Alexa-647 was excited using a Helium-Neon 633 nm laser line at 1.0% laser transmission. Laser excitation was focused into the sample by using a 40x C-Apochromat NA 1.2 water immersion objective. Correction collar was adjusted in the objective to 0.17 and room temperature. To prevent deformation of the PSF due to glycerol in the solution, the excitation volume was focused 50 µm above the glass by performing a line scan using reflected light from the 488 nm laser line. For 488-nm excitation, a 488 only main beam splitter (MBS) was used and for 633 nm and dual excitation a MBS 488/561/633 was used. Emission from Alexa-488 was collected in the range of 516–596 nm using a GaAsP detector and emission from Alexa-647 in the range of 650–694 nm using a separate GaAsP detector. Pinholes for both excitations were set to 1.0 airy units and aligned along the *xy* plane using a solution of free dye or the sample itself. Count-rate binning time was set to 1 ms and the correlator binning time was set to 0.2 µs. Count rates were never greater than 500 kHz and traces showing large deviations from the average or decaying/increasing fluorescence were manually removed from the analysis. Counts per minute (CPM) values were between 1-16 kHz for all measurements to avoid optical saturation while maximizing counts above background. For single dye measurements, 10 repetitions of 10 seconds each were collected and averaged while for receptor-ligand binding experiments 50 repetitions of 30 seconds each were collected and averaged.

### Correlation traces model fitting

FCS and FCCS raw traces were fitted using the ZEN software. Correlation and cross-correlation traces were fitted using equations that modeled the diffusion, triplet, and/or blinking processes of the fluorophores and labeled proteins. For diffusion, we assumed that the fluorescent species undergo free 3D translational diffusion. To model free 3D translational diffusion, we employed the following relation

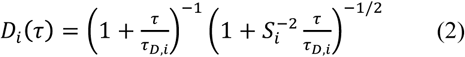

where *τ* the correlation time arising from diffusion. Equation 2 holds true in the case where there is a single fluorescent species. The suffix *i* can correspond to *g, r*, or *x*, representing the 3 channels, green, red, and cross-correlation. Interchangeable, we sometimes use for clarity the labels *R, L*, or *X* designating the green channel for the receptor labeled with Alexa-488, the red channel for the ligand labeled with Alexa-647, and the cross-correlation channel. Triplet state transitions were fitted using the following equation

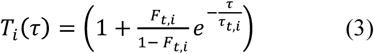

where *F*_*t,i*_ is the fraction of fluorescent species in the triplet state and *τ*_*t,i*_ is the triplet state relaxation time of the fluorophore. For Alexa-488, *τ*_*t,i*_ was set to a constant value of 4 µs and for Alexa-647, *τ*_*t,i*_ was set to a constant value of 7 µs. Equation 3 is normalized which means that the calculated number of particles excludes fluorophores present in the triplet state. Blinking state transitions were only observed when the Alexa-647 labeled chemokines were added to CCR5-SNAP-488. To model blinking state transitions, we employed the following equation,

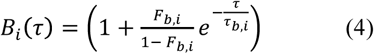

where *B*_*t,i*_ is the fraction of fluorescent species in the blinking state and *τ*_*b,i*_ is the blinking state relaxation time of the fluorophore. For Alexa-647, *B*_*t,i*_ and *τ*_*b,i*_ were fitted as free parameters.

Autocorrelation and cross-correlation traces were analyzed from 2 µs to 10 s to remove after pulsing from the detectors. For single diffusing Alexa-488 and CCR5-SNAP-488, we modeled the correlation traces using the following equation

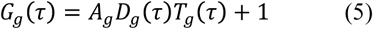

where *A*_*g*_ is the correlation amplitude for when *G*_*g*_(*τ*)= 0, *D*_*g*_*(τ*) is the diffusion correlation given by equation 2, *T*_*g*_(*τ*) is the triplet correlation given by equation 3, and 1 is an offset. *A*_*g*_, the correlation amplitude, is given by

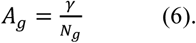

For single diffusing Alexa-647 and Alexa-647 labeled chemokines, the correlation traces were modeled using the following equation

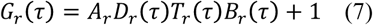

where *A*_*r*_ is the correlation amplitude for when *G*_*g*_(*τ*)= 0and *B*_*r*_ (*τ*) is the blinking correlation given by equation 4. In the absence of CCR5-SNAP-488, Alexa-647 blinking is not observed and the correlation amplitude was fitted using equation 5. *A*_*r*_, the correlation amplitude, is given by

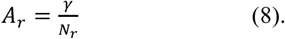

To fit the cross-correlation traces, we employed the following equation

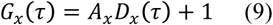

where the cross-correlation amplitude, *A*_*X*_, is given by

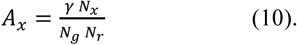

For all 3 channels, *γ* is set to 0.35 and the structural parameter to 8. To derive errors for each measurement, the total repetitions were divided into 3 independent sets of measurements and each set was averaged and analyzed using the equations above. From these 3 averages, the standard deviation was calculated for the number of particles. In cases where *τ*_*D*_*complex* deviated significantly from previously measured values, it was fixed to 550 µs so that the fit would converge. From the calculated number of particles, we derived concentrations using equations 14-16 below, which were used for global analysis of ligand binding.

### Confocal volume determinatio

Alexa-488 and Alexa-647 solutions were diluted in Buffer N at various concentrations from 25 nM to 0.8 nM. FCS measurements were acquired as described above. The number of particles derived from the fits of the correlation traces was plotted as a function of concentration (Supplementary Figure S3). The concentration of the Alexa-488 and Alexa-647 stock solutions used for the dilutions (nominally 10 µM) was determined by UV-Vis. The UV-Vis derived concentration was used instead of the nominal fluorophore concentrations for calculating the concentrations of the calibration solutions. To determine the confocal volume for the cross-correlation (CC) channel, we employed a dual-labeled oligonucleotide 40 base pairs long with Alexa-488 and Alexa-647 at both ends to minimize FRET based similar to the original design from Schwille et al., who used rhodamine green and Cy5 as fluorophores.(20)

The oligos were synthesized by IDT. The forward sequence for the oligonucleotide is 5’-[AminoC6Alexa488]GCCGT CTCTGACTGCTGATGACTACTATCGTATAGTGCGG[BioTEG-Q]-3’ and the sequence for the reverse oligonucleotide is 5’-[AminoC6Alexa647]CCGCACTATACGATAGTAGTCATCAGCAGTCAGAGACGGC-3’. Single strand oligonucleotides were re-suspended in 10 mM Tris pH 8.0, 50 mM NaCl, 1 mM EDTA to a final concentration of 1 µM in 100 µL of buffer. Samples were added to a water bath at 94 °C in a Dewar flask and allowed to anneal until the temperature in the water bath reached less than 40 °C. Oligos were then placed at room-temperature while a C4 column was equilibrated with 0.1 M ammonium acetate at pH 6.6. Oligos were loaded into the column and eluted with a 0–50% (v/v) gradient of acetonitrile in water. HPLC fractions were analyzed by UV-Vis absorbance for both Alexa-488 and Alexa-647. The peak fraction was aliquoted into 50 µL aliquots and stored at −20 °C for long-term storage. The final concentration of the oligo was 110 nM. Stock solutions were employed to dilute the oligonucleotide as done for the free dyes.

We determined the confocal volumes (*V*_*g*_, *V*_*r*_, or *V*_*x*_) using the observed number of particles (*N*_*g*_, *N*_*r*_, or *N*_*x*_) from the linear fits of the experimentally observed number of particles as a function of concentration (*C*_*g*_, *C*_*r*_, or *C*_*gr*_) (Eqs. 11-13). The slope of the linear fit divided by Avogadro’s number yields the confocal volumes (see Supplementary Figure S3).

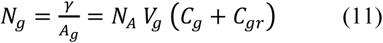

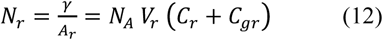

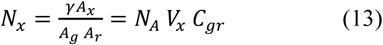

The following relations for the green and red channels relate the sample concentrations to the correlation amplitudes and the number of particles derived from the amplitudes:

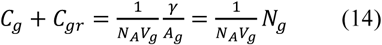

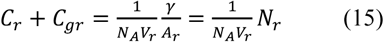

Where *C*_*g*_ and *C*_*r*_ are the concentrations of the green and red particles respectively, and *C*_*gr*_ is the concentration of the double-labeled species. In the case where there is only a single fluorescent species or no binding, the *C*_*gr*_ term vanishes to 0. *NA* is Avogadro’s number, *V*_*g*_ and *V*_*r*_ are the confocal volumes of the green and red channels respectively, *A*_*g*_ and *A*_*r*_ are the correlation amplitudes for the green and red channels respectively, *N*_*g*_ and *N*_*r*_ are the number of particles of the green and red particles respectively, and *γ* is a correction factor to account for the fact that the confocal volume deviates from the 3D Gaussian approximation.

For the cross-correlation, we employed the following relation for the concentration,

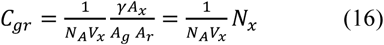

where *V*_*x*_ is the cross-correlation confocal volume, *A*_*x*_ is the cross-correlation amplitude, and *N*_*x*_ is the number of particles that is double-labeled.

The confocal volume is related to its dimensions by the following relations for the green and red channels:

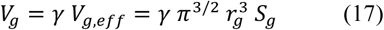

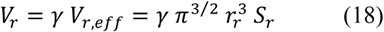

*V*_*g,eff*_ and *V*_*g,eff*_ are the effective confocal volumes of the green and red channels respectively, *r*_*g*_ and *r*_*r*_ are the radii of the confocal volumes along the *xy* plane, and *S*_*g*_ and *S*_*r*_ are the structural parameters of the green channel and red channels respectively. The structural parameter is defined 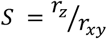 where *r*_*z*_ and *r*_*xy*_ are the radii along the *z* and *xy* planes respectively for each channel.

Since the cross-correlation confocal volume is defined as the overlap between the green and red confocal volumes, then the equation describing the cross-correlation confocal volume is given by

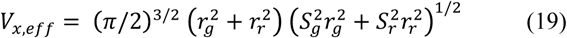

Equation 19 is only valid for the case in which there are no chromatic aberrations in the system. The radii of the confocal volumes relate the diffusion coefficients to the observable diffusion times:

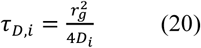

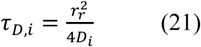

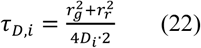

where *τ*_*D,i*_ is the diffusion time of species *i* and *D* is the diffusion coefficient of the same species.

### Crosstalk determination

Cross-talk from the green channel to the red channel was determined using the protocol by Bacia & Schwille (2007).(12) Briefly, Alexa-488 (25 nM) in Buffer N was excited using 488-nm laser line and the count rates were recorded using a GaAsP detector. The same solution was excited with the same laser line but instead count rates were recorded on a separate GaAsP detector, which is used for the red channels. Count rates from the two measurements were used to calculate the bleed-through ratio:

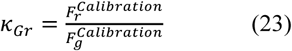

Under our experimental conditions, the bleed-through ratio is 0.0072. In a sample containing dual-labeled oligonucleotide or CCR5-SNAP-488 and labeled chemokine, the ratio of the measured green and red count rates was taken and then multiplied with the bleed-through ratio calculated previously to obtain the cross-correlation amplitude relative to the green auto-correlation as expected from cross-talk only.

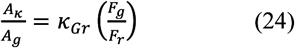

We obtained a value of 0.0033, which is less than 0.4% of the observed relative cross-correlation not corrected for crosstalk obtained by equation 25:

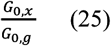

Therefore, crosstalk plays only a minor role and can be neglected unless otherwise noted.

### Global Fitting Analysis of Saturation and Competition Binding Curves

For ligand binding assays, we assumed that the chemokines recognize a single binding site on CCR5. Based on this, we then set the concentration of complexes, *C*_*gr*_, equal to

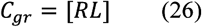

where [*RL*] is the total concentration of receptor-ligand complexes. In the green channel, the calculated number of particles arise from both receptor and receptor-ligand complexes. Therefore,

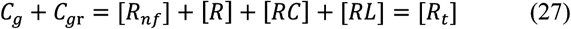

where [*R*_*nf*_] is equal to the concentration of non-functional receptor species, [*R*] is the concentration of ligand-free receptor, [*RC*] is the concentration of receptor-competitor complex, [*RL*] is the concentration of receptor-ligand complex, and [*R*_*t*_] is the total concentration of receptor. To define the fractional occupancy, we employ the following relation

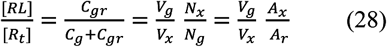

which shows that fractional occupancy can be derived from the ratio of 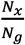. Alternatively, the fractional occupancy is can also be derived from the ratio 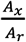. Since we have non-functional receptor species, we define [*R*_*nf*_] as follows

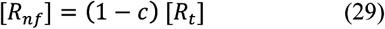

where *c* is a constant to define the fraction of receptor that is functional. Therefore, we can state that

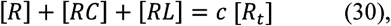

the fractional occupancy for the ligand-bound receptor becomes

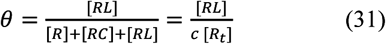

and for the competitor-bound receptor

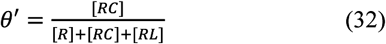

In the red channel, we observe fluorescence from the free ligand and the receptor-ligand complex, therefore the concentrations of each are given by

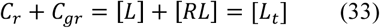

where [*L*_*t*_] is the total concentration of ligand.

For a single binding site, the equilibrium dissociation and inhibition constants are defined as follows

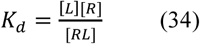

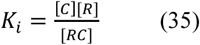

Where *K*_*d*_ is the equilibrium dissociation constant, and *K*_*i*_ is the equilibrium inhibition constant. *C* and *L* are the free concentrations of competitor and ligand, respectively. *R* is the free concentration of receptor. *RL* is the concentration of receptor-ligand complex. *RC* is the concentration of receptor-competitor.

Using the expressions above, we derived the following equations for fractional occupancy in terms of free ligand, free competitor, *K*_*d*_ and *K*_*i*_

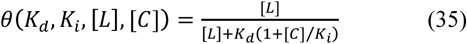

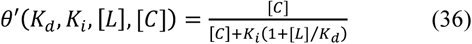

Since the FCCS observables for concentrations are the total number of species for each channel, we developed an iterative algorithm to calculate the concentrations of free ligand and competitor. With this algorithm, the concentrations of free ligand, free competitor, equilibrium dissociation constant, and equilibrium inhibition constant, are fitted as free parameters on the saturation and competition binding isotherms. The algorithm is repeated multiple times until the fitted values match the values given by the fractional occupancy calculated from FCCS measurements. In other words, we minimize *k* in the reduced chi-square function (eq. 37) shown below

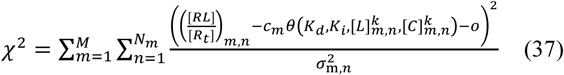

Where 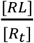 is derived from FCCS and FCS measurements, the suffix *m* is to represent each individual experiment, and the suffix *n* is to represent each data point per experiment. The variance, 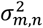, is set equal to 1. The variable, *o*, corresponds to an offset to include a small crosstalk correction, which accounts for the fact that the competition binding isotherms don’t fully reach 0 under saturating concentrations of competitor. The value of *o* was fixed to 0.01.

We account for ligand depletion by an iterative algorithm. We initialize the algorithm by setting the free ligand and competitor concentrations to their total concentration:

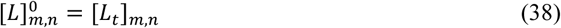

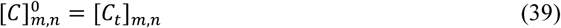

With these initial free concentrations, we solve the binding equilibrium to calculate the concentration of ligand-receptor and competitor-receptor complexes. Next, we re-estimate the free ligand and receptor concentrations as the difference of total and bound concentrations:

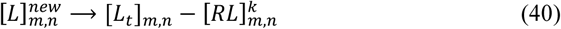

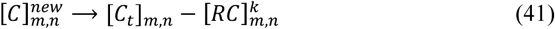

Instead of using these new values for the free concentrations directly in the next iteration, we update the free concentrations for the next iteration more gradually to avoid an oscillatory behavior of the algorithm:

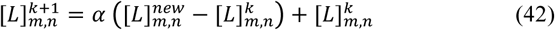

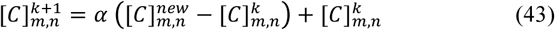

where *k* ⟶ *k* +1 is the update of the iteration counter. The parameter α can be used to tune the algorithm. We use α *=* 0*.25* with 20 iterations. Using equations 40–43 for the ligand depletion, we minimize eq. 37 in each iteration to yield a converged set of global fitting parameters that optimally fit the data.

Global analysis with non-linear least square fitting of the binding isotherms yielded parameters that describe the complete data set. Saturation and competition binding isotherms were normalized and binding isotherms were plotted as a 2D function (ρ, *L*) or a 3D surface (ρ, *C, L*) for saturation and competition binding, respectively. To determine the errors associated with each affinity, we performed a statistical bootstrapping error analysis by random data resampling with replacement. We resampled the data one hundred times and calculated the means and standard deviations of all model parameters. In the tables, we report the model parameters determined from the original fit together with the standard deviations of these parameters from the bootstrap analysis. The means from the bootstrap analysis were always within the error bounds of the original solution.

## RESULTS

We designed a codon-optimized human CCR5 synthetic gene with a C-terminal SNAP-tag fusion to facilitate high-level expression and covalent fluorescent labeling. We positioned the SNAP-tag on the intracellular C-terminal tail of CCR5 to minimize FRET with the fluorescently-labeled RANTES analog ligands that bind at the extracellular surface. Although this construct expresses at higher levels than WT CCR5 in HEK293T cells (Supplementary Fig. S4), we found that the expressed receptor was subject to proteolytic degradation at the N-and C-terminal tails. In order to obtain homogeneous full-length receptor for quantitative fluorescence measurements, we modified the CCR5 construct with a cleavable signal peptide from the serotonin 5-HT_3A_ receptor to enhance proper membrane insertion of the receptor with added N-terminal FLAG-tag to allow tandem affinity purification (Fig. 1a and Fig. 2).(21) Two OLLAS mAb epitope tags (*Escherichia coli* OmpF Linker and mouse Langerin fusion Sequence)(22) and the Strep III tag(23) were included for future applications, such as receptor immobilization on functionalized surfaces and alternative purification procedures (Fig. 1a). We analyzed expression of this new construct, referred to as simply CCR5-SNAP, by dual-color near infrared (NIR) immunoblot analysis and observed that the full-length receptor migrated as a 1D4/FLAG dual stained (red&green) band with an apparent molecular mass of approximately 70 kDa (Fig. 1b).

**Figure 1.**
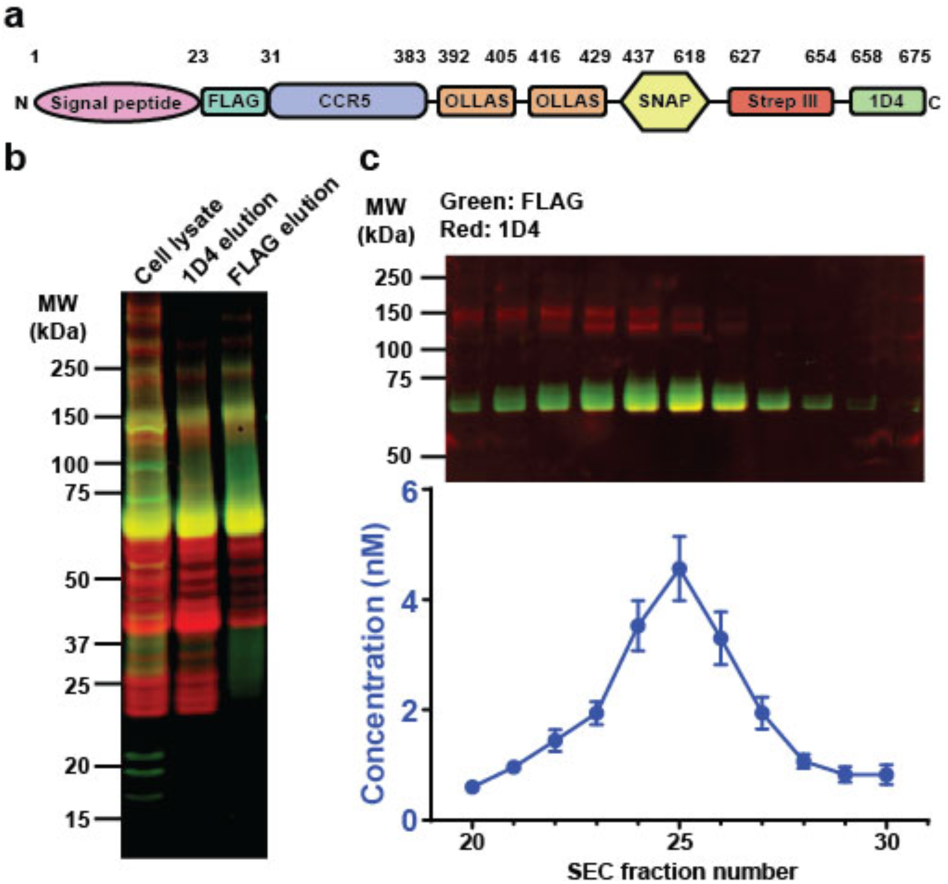
Expression, labeling and purification of CCR5-SNAP. (a) CCR5-SNAP construct schematic showing the receptor (blue) fused to the signal peptide (violet) for proper receptor insertion into the membrane, FLAG (cyan) and 1D4 (green) epitopes for tandem affinity purification, double OLLAS (orange) and Strep-III (red) epitopes for optional surface immobilization and purification, and SNAP-tag (yellow) for fluorescent labeling. (b) Reducing SDS-PAGE and NIR-immunoblot of cell lysate, 1D4 and FLAG elutions from the 1D4/FLAG tandem affinity purification. Full-length CCR5-SNAP (∼70 kDa, yellow band) was detected using antibodies against the 1D4 (red) and FLAG (green) epitopes. (c) Representative reducing SDS-PAGE and NIR-immunoblot from one CCR5-SNAP-488 SEC purification. Average values from FCS-derived concentrations for SEC fractions 20 to 30 are from five independent CCR5-SNAP-488 SEC purifications. Error bars are ± S.E.M. The purification protocol is shown schematically in Fig. 2.

**Figure 2.**
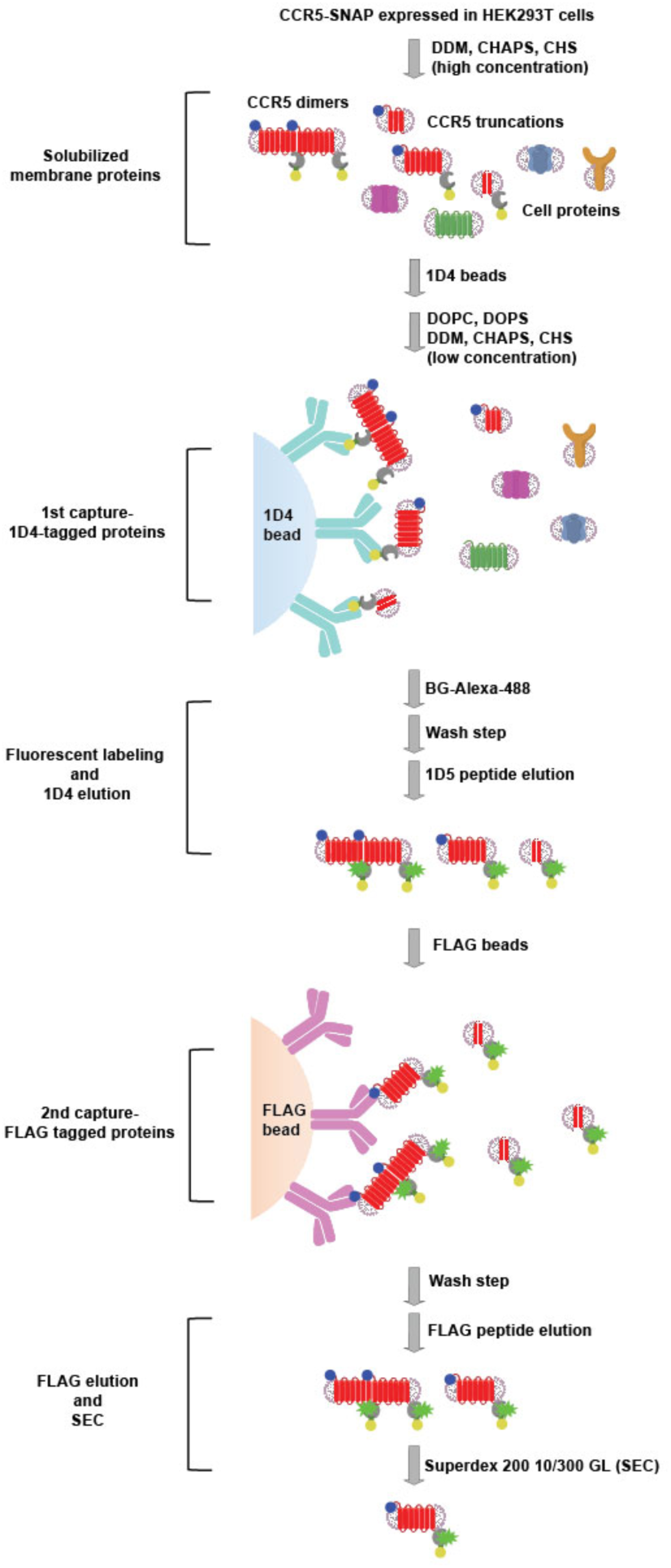
CCR5-SNAP purification scheme. Schematic displaying the combined tandem affinity and size exclusion purification strategy to yield full-length, monomeric CCR5-SNAP-488 from mammalian cells. HEK293T cells transiently expressing CCR5-SNAP are lysed in buffered solution containing DDM, CHAPS, and CHS (see Materials and Methods). Cell lysate, which contained cellular proteins, full-length CCR5-SNAP, and CCR5-SNAP truncations, was added to 1D4 resin. Lipids DOPC and DOPS were added to the buffered solution and then CCR5-SNAP was labeled with the SNAP substrate BG-Alexa-488. Excess fluorophore was washed away and the labeled receptor was eluted using excess 1D5 peptide. The 1D4 eluate was added to FLAG beads to bind full-length receptors and remove receptor truncations. FLAG resin was washed and full-length CCR5-SNAP was eluted using excess FLAG peptide. FLAG elution was applied to a Superdex 200 10/300 GL column to purify monomeric CCR5-SNAP from receptor aggregates.

We then employed a tandem affinity purification strategy to isolate full-length CCR5-SNAP from receptor truncation products. Transiently transfected HEK293T cells expressing CCR5-SNAP were lysed with a buffer containing DDM, CHAPS, and CHS. The solubilized receptor was immobilized onto the 1D4-Sepharose immunoaffinity matrix (Fig. 2). The detergent concentration was lowered and lipids (DOPC and DOPS) were added to enhance receptor stability(24, 25) and CCR5-SNAP was fluorescently labeled on-resin with benzyl guanine-Alexa-488 (BG-Alexa-488), washed several times to remove excess dye, and then eluted using 1D4 nonapeptide. The first step of the purification removes cellular components and any C-terminal receptor truncations, but not N-terminal truncations (Fig. 1b). To purify full-length receptor from these truncations, we performed a second immunoaffinity purification step using anti-FLAG M2 agarose, which removed N-terminal truncations to yield the desired full-length receptor (Fig. 1b). NIR-immunoblot analysis of 1D4/FLAG-purified CCR5-SNAP-488 shows the presence of minor bands corresponding to SDS-resistant CCR5-SNAP dimers (∼150 kDa) and oligomers (∼250 kDa) (Fig. 1b).

Since oligomers complicate the ligand-binding analysis, we used size-exclusion chromatography (SEC) to purify receptor monomers from undesired oligomers. We determined the concentration of CCR5-SNAP-488 in the SEC fractions by FCS (Fig. 1c and Supplementary Fig. S5) as the concentrations were too low for reliable quantification by other methods including 280-nm absorbance. The peak CCR5-SNAP fraction had an average concentration of (4.5 ± 0.6) nM. Because we observed a single peak in the FCS-derived chromatogram, we also analyzed the SEC fractions by NIR-immunoblot to evaluate the separation of monomeric CCR5-SNAP-488 from oligomers. Monomeric CCR5-SNAP-488 eluted as a single species while it co-eluted with receptor oligomers in earlier fractions (Fig. 1c and Supplementary Fig. S5). Monomeric full-length receptor was employed for all FCCS ligand-binding measurements.

We characterized expressed CCR5-SNAP in cell-based functional assays to determine whether the engineered tags interfere with receptor function. Since CCR5-SNAP expressed at higher levels than wild-type 1D4-tagged CCR5 (CCR5), we optimized the gene dosage of CCR5-SNAP to obtain similar surface expression as CCR5 to accurately compare their function and pharmacology. We quantified CCR5-SNAP and CCR5 cell surface expression by flow cytometry using the anti-CCR5 antibody 2D7 conjugated to PE (Supplementary Fig. S6a). CCR5-SNAP and CCR5 expressed at similar levels when HEK293T cells were transfected with 2.0 µg and 0.75 µg of plasmid DNA, respectively. To validate the results, we repeated the experiment using the T21/8-PE antibody, which recognizes a different epitope than 2D7. CCR5-SNAP and CCR5 expressed at similar levels, in agreement with the results obtained with the 2D7-PE antibody (Supplementary Fig. S6a). As a further control, we quantified CCR5-SNAP expression using the FLAG-PE antibody and we observed fluorescence only from CCR5-SNAP expressing cells, but not from CCR5 expressing cells (Supplementary Fig. S6a). We also performed immunofluorescence TIRF imaging on CCR5-SNAP or CCR5 expressing HEK293T cells and we observed similar receptor expression patterns between CCR5-SNAP and CCR5 (Supplementary Fig. S6d).

We next measured Gαi-protein activation-dependent inhibition of forskolin-stimulated cyclic AMP (cAMP) levels in HEK293T cells transfected with CCR5-SNAP and CCR5 in response to various chemokine concentrations using the RLuc3-EPAC-GFP10 BRET^2^ reporter.(14, 16) We fitted the dose-response curves for WT CCR5 and CCR5-SNAP as a function of chemokine concentrations using a three-parameter logistic equation to derive EC_50_ and E_max_ values (Supplementary Fig. S6b). Both CCR5-SNAP and CCR5 inhibited cAMP production for all the tested chemokines with similar efficacies and potencies (Supplementary Table S5).

We also measured calcium mobilization after chemokine stimulus in CCR5-SNAP and CCR5 expressing HEK293T cells using the FLIPR calcium 6 dye and generated dose-response curves (Supplementary Fig. S6c). 5P12 and 5P14 did not induce calcium flux on CCR5-SNAP or CCR5 expressing cells in agreement with previous literature reports.(9) For the remaining chemokines tested, we did not observe any meaningful differences in efficacy or potency between CCR5-SNAP and CCR5 (Supplementary Table S6). We conclude from these functional control experiments that the SNAP-tag does not seem to interfere with biological function of the receptor.

In FCS and FCCS experiments, autocorrelation and cross-correlation amplitudes, denoted as *A*, are inversely proportional to the concentration of the fluorescent species (Materials and Methods). To derive concentrations from *A*, we determined the size of the confocal volumes for 488-nm excitation (0.18 ± 0.02 fL), 633-nm excitation (0.35 ± 0.03 fL), and their volume overlap (0.22 ± 0.03 fL) using serial dilutions of Alexa-488, Alexa-647, and a 40-base pair Alexa-488/Alexa-647-dual-labeled oligonucleotide duplex, respectively (Supplementary Fig. S3).(20, 26)

To illustrate the changes of auto-and cross-correlation functions in a ligand-binding experiment, we modeled the correlation function amplitudes as a function of titrating ligand concentration for receptor, ligand and receptor-ligand complex at equilibrium dissociation constants of *K*_*d*_ = 5 nM and *K*_*d*_ = 0.5 nM (Fig. 3a). *A*_*R*_, the receptor correlation amplitude, remains constant while *A*_*L*_, the ligand correlation amplitude, grows hyperbolically with decreasing concentration. In contrast, *A*_*X*_, the receptor-ligand complex fluorescence cross-correlation amplitude, plateaus at low ligand concentrations and asymptotically goes to zero at high ligand concentrations. Higher ligand affinities yield larger *A*_*X*_ amplitudes as compared with lower ligand affinities because more receptor-ligand complexes are present. Assuming equal confocal volumes for all channels, the ratio *A*_*X*_/*A*_*L*_ is equal to the fractional receptor occupancy and a plot of *A*_*X*_/*A*_*L*_ as a function of ligand concentration yields a saturation binding isotherm (Fig. 3a). We also simulated two different competition-binding cases where ligand and receptor concentrations were kept constant while an unlabeled competitor concentration was titrated (Fig. 3b). For these simulations, we assumed the following ligand and competitor affinities: a) *K*_*d*_ = 0.5 nM and *K*_*i*_ = 0.5 nM and b) *K*_*d*_ = 0.5 nM and *K*_*i*_ = 5 nM (Fig. 3b). *A*_*R*_ and *A*_*L*_ remain constant while *A*_*X*_ decreases with increasing competitor concentration. Similarly, we can derive competition binding isotherms by plotting *A*_*X*_/*A*_*L*_ as a function of competitor concentration (Fig. 3b). Thus, we show that fractional occupancy can be derived from correlation amplitudes to derive saturation and competition binding isotherms.

**Figure 3.**
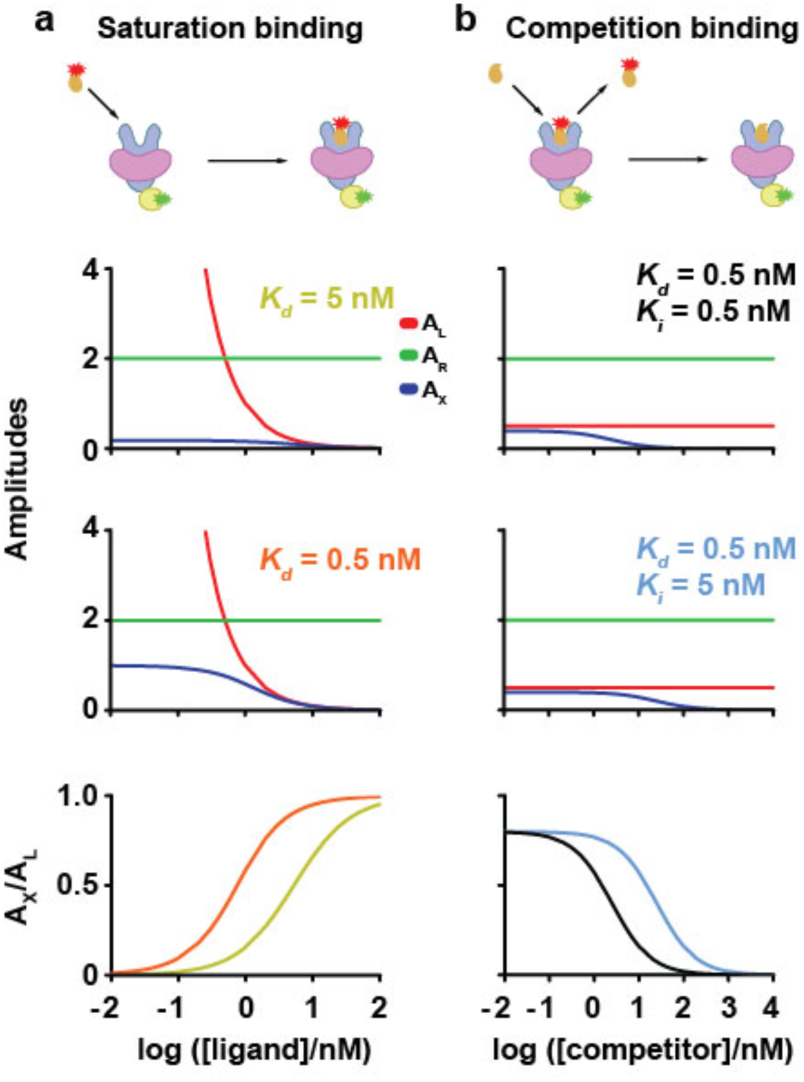
Modeling binding parameters from FCCS measurements. Fractional receptor occupancy can be derived from correlation amplitudes to derive saturation (column a) and competition (column b) binding isotherms. Cartoons show the binding interactions analyzed by FCCS. Fluorescent ligand (orange with red star) recognizes a lipid-embedded (violet) membrane receptor (blue). The receptor is fused to a functional tag (yellow) labeled with a fluorophore (green). The amplitudes are modeled as a function of titrating ligand concentration for receptor, ligand and receptor-ligand complex. For saturation binding (column a), correlation amplitude plots are shown for receptor (green, *A*_*R*_), ligand (red, *A*_*L*_), and complex (blue, *A*_*X*_) as a function of fluorescent ligand concentrations where ligand dissociation constants were set to 5 nM (yellow) or 0.5 nM (orange). Ligand concentration varied from 0.01 to 100 nM and receptor concentration was kept constant at 0.5 nM. For competition binding analysis (column b), competitor concentration varied from 0.01 to 10,000 nM and ligand and receptor concentrations were kept constant at 2 and 0.5 nM, respectively. Competition binding isotherms can be derived by plotting *A*_*X*_/*A*_*L*_ as a function of competitor concentration. Two different competition-binding cases were simulated where ligand and receptor concentrations were kept constant while an unlabeled competitor concentration was titrated (column b). The ligand and competitor affinities were assumed to be *K*_*d*_ = 0.5 nM, *K*_*i*_ = 0.5 nM (black) or *K*_*d*_ = 0.5 nM, *K*_*i*_ = 5 nM (cyan). *A*_*R*_ and *A*_*L*_ remain constant while *A*_*X*_ decreases with increasing competitor concentration.

The simulations assume that there is no crosstalk from the green channel to the red channel. Crosstalk from the green channel to the red channel increases *A*_*X*_, leading to an overestimation of the concentration of receptor-ligand complexes. Given this, we quantified the crosstalk contribution to *A*_*X*_ using the method by Bacia & Schwille (2007)(12) and we measured negligible crosstalk contribution to *A*_*X*_ corresponding to an error in receptor occupancy of less than one percent.

We then performed saturation-binding experiments with synthetic RANTES analogs site-specifically and covalently labeled with Alexa-647 (Supplementary Fig. S2) and purified, monomeric CCR5-SNAP-488 by FCCS to determine their equilibrium dissociation constants. Representative autocorrelation and cross-correlation traces and fits are shown in Fig. 4 and Supplementary Fig. S7. We also performed homologous and heterologous competition binding FCCS experiments using Alexa-647-labeled 5P12 (5P12-647) and 6P4 (6P4-647) with unlabeled 5P12 and 6P4 (Fig. 5 and Supplementary Fig. S8). Before further analyzing the data, we plotted the experimental *G*(0) values for the ligand, receptor and ligand-receptor complex as a function of nominal ligand concentration for both the saturation and competition binding experiments and we observed non-random variations in *G*(0) for all three components (Supplementary Fig. S7d and S8d) that correlate with non-random variations in the diffusion time (Supplementary Fig. S9) most likely due to changes in the point-spread-function across the glass-bottom microtiter plate. A more complete discussion of variations in the diffusion time and molecular brightness is presented in the supplement (Supplementary Text and Figs. S10–S12).

**Figure 4.**
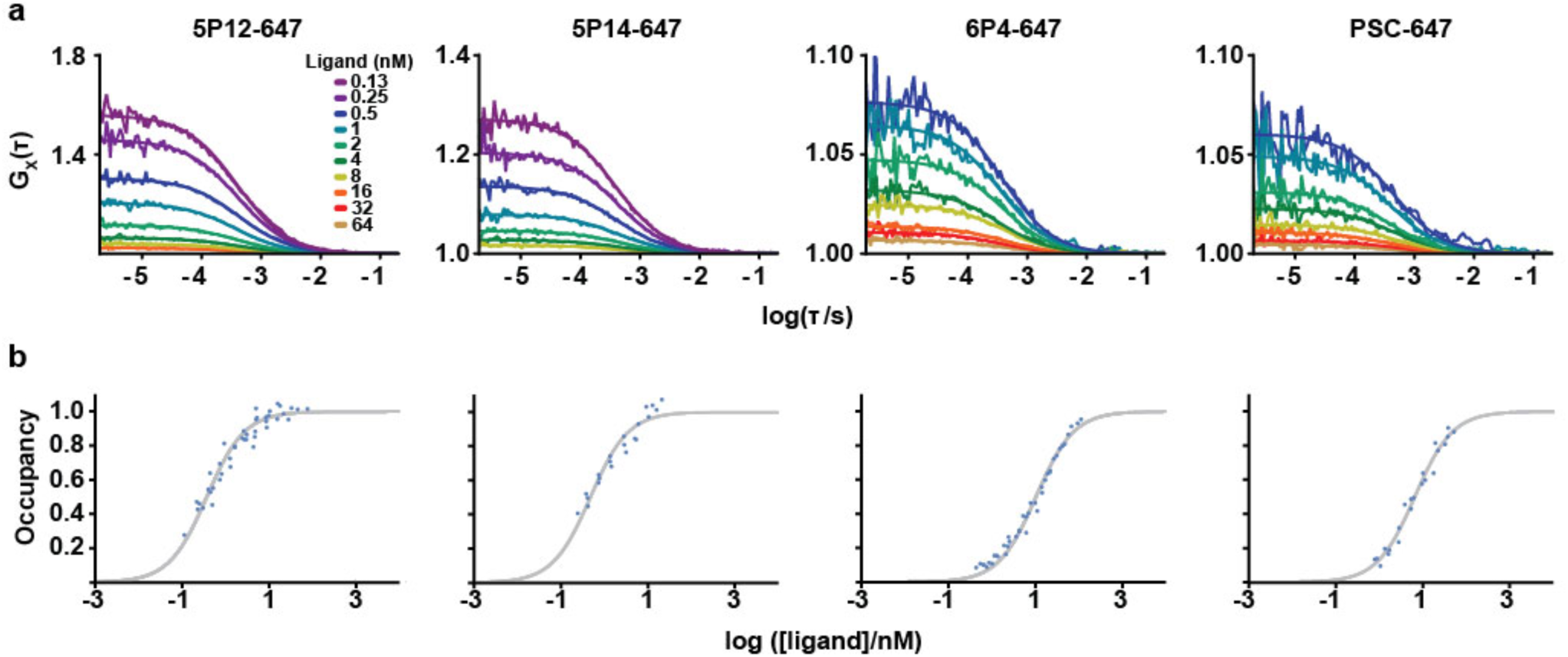
Saturation binding of Alexa-647 labeled RANTES analogs to CCR5-SNAP-488 by FCCS. (a) Cross-correlation traces and fits from one representative experiment of 5P12-647, 5P14-647, 6P4-647 and PSC-647 binding to CCR5-SNAP-488 at various ligand concentrations. Cross-correlation traces were fitted to one translational component with 3D diffusion. (b) Saturation binding isotherms for 5P12-647, 5P14-647, 6P4-647, and PSC-647 binding to CCR5-SNAP-488. Ligand binding fits were performed using data from at least three independent experiments and assuming 25% of CCR5-SNAP-488 is functional.

**Figure 5.**
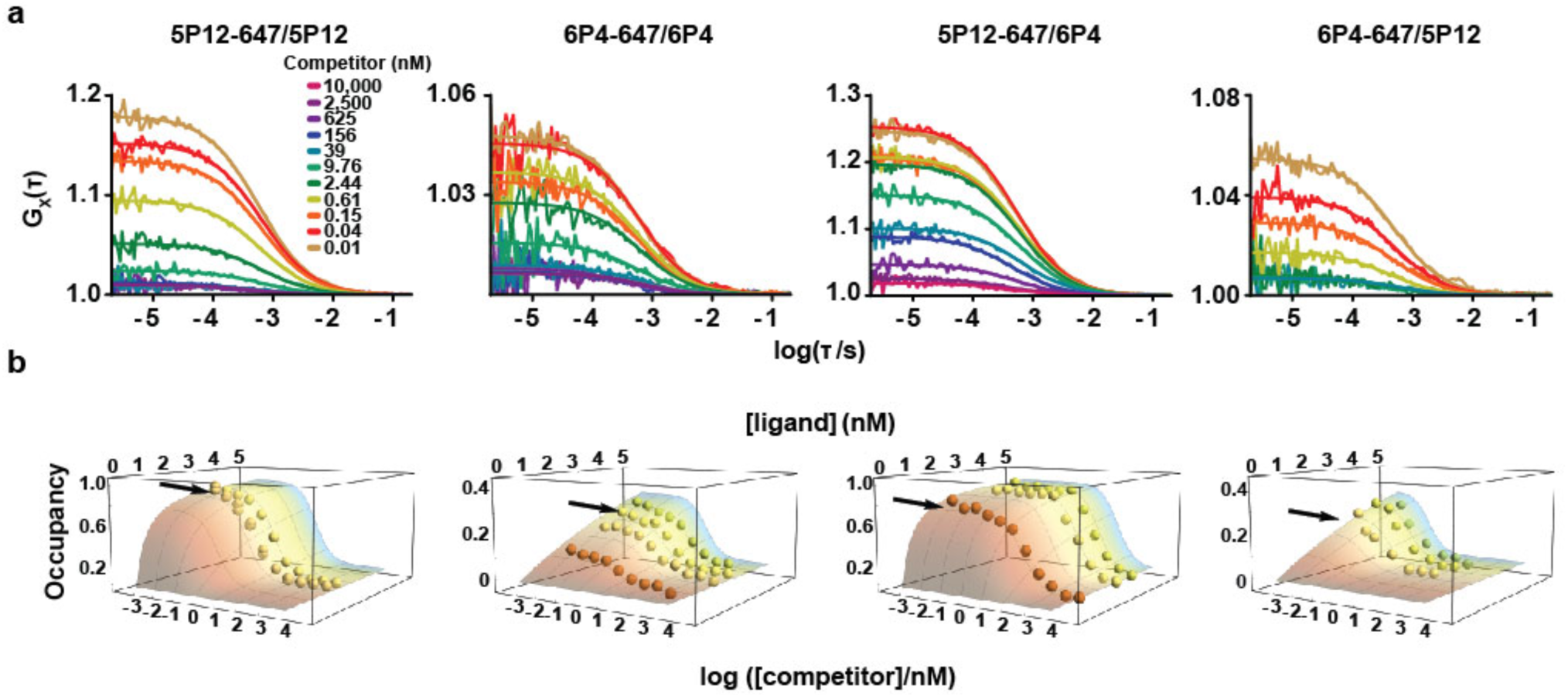
Competition binding of Alexa-647 labeled 5P12 and 6P4 with non-labeled chemokines by FCCS. (a) Cross-correlation traces and fits from one representative experiment of homologous and heterologous competition binding with 5P12-647 and 6P4-647 at various competitor and ligand concentrations. Cross-correlation traces were fitted to one translational component with 3D diffusion. (b) Homologous and heterologous competition binding isotherms for 5P12-647 and 6P4-647. Competition binding data were plotted as 3D surfaces to show the effect of ligand variability on the normalized occupancy. Arrows point to the representative experiments shown in (a). Ligand binding fits were performed using data from at least three independent experiments and with 25% functional CCR5-SNAP-488.

Since the fractional occupancy derived from saturation binding experiments was less than 100% for each of the RANTES analogs tested, we postulated that some fraction of CCR5-SNAP-488 becomes inactive during purification. We accounted for this inactive fraction and introduced a free parameter, *c*, in the ligand-binding model to specify the active receptor fraction for each experimental data set. We performed global fitting of the binding curves using an iterative algorithm that accounts for ligand depletion by reducing the chi-square function defined in the Materials and Methods. Fig. 4b shows saturation binding isotherms and fits for the RANTES analogs. From these fits, we calculated *K*_*d*_ = (0.37 ± 0.06) nM for 5P12-647 and *K*_*d*_ = (0.48 ± 0.08) nM for 5P14-647 (Table 1). In contrast, we derived *K*_*d*_ = (10.4 ± 1.2) nM for 6P4-647 and *K*_*d*_ = (6.6 ± 0.8) nM for PSC-647 (Table 1). Based on these results, we conclude that if 6P4-and PSC-647 are already high affinity binders to CCR5-SNAP-488 given their nanomolar affinity, then 5P12-and 5P14-647 should be considered ultra-high affinity binders.

**Table 1.** FCCS derived binding affinities of RANTES analogs at CCR5-SNAP-488. Equilibrium dissociation (*K*_*d*_) and inhibition (*K*_*i*_) constants for the RANTES analogs binding to the CCR5-SNAP-488 derived from saturation and competition binding. Errors were derived using global analysis with non-linear least square fitting of the binding isotherms with bootstrapping.

| Chemokine | $K_d$ / nM |
| --- | --- |
| 5P12-647 | $0.368 \pm 0.064$ |
| 5P14-647 | $0.475 \pm 0.082$ |
| 6P4-647 | $10.4 \pm 1.2$ |
| PSC-647 | $6.57 \pm 0.81$ |

Table 1. FCCS derived binding affinities of RANTES analogs at CCR5-SNAP-488.
| Labeled/Unlabeled | $K_i$ / nM |
| --- | --- |
| 5P12-647/5P12 | $0.141 \pm 0.033$ |
| 5P12-647/6P4 | $11.8 \pm 2.0$ |
| 6P4-647/6P4 | $3.49 \pm 0.77$ |
| 6P4-647/5P12 | $0.047 \pm 0.099$ |

The RANTES analogs are labeled with Alexa-647, which contains negatively charged sulfonate groups that might perturb the binding of the positively charged chemokines to the receptor. We therefore performed homologous competition binding experiments with 5P12-647 and 6P4-647 with titrating concentrations of non-labeled 5P12 and 6P4, respectively. We performed the same global fitting analysis as with the saturation binding measurements and we employed the variation in labeled ligand to plot fractional occupancy as a function of both ligand and competitor. Fig. 5b shows the 3D competition-binding surface plots for 5P12 and 6P4 homologous competition binding. We derived *K*_*i*_ = (0.14 ± 0.03) nM for 5P12, and *K*_*i*_ = (3.5 ± 0.8) nM for 6P4 (Table 1). The calculated *K*_*i*_ values for 5P12 and 6P4 are approximately three-fold lower as compared to the *K*_*d*_ values of the labeled ligands calculated for saturation binding, indicating that Alexa-647 slightly perturbs chemokine binding to CCR5-SNAP-488, likely as a result of the electrostatic repulsion of the negatively charged fluorophore and the receptor in the negatively charged mixed micelle.

## DISCUSSION

FCS is a single-molecule-sensitive method in which fluorescence fluctuations from molecules diffusing through a sub-femtoliter volume of focused excitation are analyzed.(27) An autocorrelation analysis is performed on the fluorescence fluctuations to derive physical parameters such as diffusion coefficients, triplet state fractions and concentrations. Because of the small size of the diffraction limited volume in FCS measurements, even sub-nanomolar concentrations of ligand can be studied. In contrast, techniques such as surface plasmon resonance (SPR) and isothermal calorimetry generally require at least micromolar concentrations of bound ligand in the detection volume to measure ligand-binding affinities, which complicates determination of nanomolar affinities. Another advantage of FCS is that binding interactions are observed in solution, overcoming immobilization artifacts present in SPR and in some other imaging techniques.(26) However, FCS alone cannot discern binding between two species that have similar molecular weights. In order to separate the free and bound species in the FCS autocorrelation curve, a fluorescently-labeled ligand must undergo at least an eight-fold change in molecular mass upon binding to its receptor. This limitation can be surmounted if the receptor and ligand can be labeled with different fluorophores emitting at distinct wavelengths so that two autocorrelation curves can be measured simultaneously to determine a so-called cross-correlation function using FCCS. We used the FCCS approach to discover and study ultra-high affinity interactions between CCR5 and a series of RANTES analogs of biological significance.

In order to enable quantitative measurements of ligand affinities for CCR5 using FCCS, we purified to near homogeneity a monomeric fluorescently-labeled CCR5-SNAP-488 from receptor oligomers, aggregates, and truncation products originating from both N- and C-terminal degradation. The purified receptor was quantified in SEC fractions using FCS. Previously, Nisius *et al.* purified CCR5 from insect cells and observed in the size-exclusion chromatograms two distinct peaks corresponding to equal fractions of monomeric and dimeric receptor.(28) In comparison, we only observed a very small amount of dimers, most likely due to the mammalian cell expression system and the different detergents employed, which may alter the extent of receptor oligomerization. We demonstrated that the expressed CCR5-SNAP construct signals like wild-type CCR5 in cAMP inhibition and calcium mobilization assays in response to agonist ligands, showing that the SNAP-tag and additional affinity tags do not interfere with receptor function.

Using the FCCS method, we calculated the equilibrium binding affinities for the fluorescently-labeled RANTES analogs at the purified functional CCR5-SNAP-488.

Antoine *et al.* performed similar ligand binding measurements by FCCS using several green fluorescent protein (GFP) receptor fusion proteins with fluorescently-labeled small molecules and antibodies.(29) Our approach differs in two major ways. First, we used highly purified receptor samples for ligand-binding measurements, instead of GPCRs solubilized in crude cell lysates without any purification. As such, effectors that regulate ligand binding are not removed and the measured affinities are not representative of the naked receptor. Also, we used the SNAP-tag technology to label receptors, rather than GFP as a fluorescent label fused to the target receptor. GFP is susceptible to misfolding during biosynthesis, leading to sub-stoichiometric receptor fluorescent labeling. Using the SNAP-tag, we achieved stoichiometric labeling of functional CCR5-SNAP by supplementing the reaction buffer with 1 mM DTT.

We also observed that the native chemokines RANTES and MIP1α could not displace 5P12-647 or 6P4-647 from CCR5-SNAP-488 (Supplementary Text and Figs. S13–S16). This results parallels earlier reports that PSC-biotin binding could not be displaced with RANTES or MIP-1α, but only with PSC.(30) We expected that sCD4/gp120 complex should also recognize the “naked” receptor fraction, but we could not displace either 5P12-647 or 6P4-647 in our competition binding experiments with the viral complex. In previous cell based assays, the radiolabeled sCD4/gp120 complexes in cells could be displaced by the RANTES analogs,(31) but it is unknown whether the viral complex can compete off the RANTES analogs in cells.

We demonstrated that the RANTES analogs bind with high affinity to the naked CCR5 receptor. In contrast, the native chemokines RANTES and MIP1α did not bind to the CCR5 in detergent solution. We ruled out that the tags on the CCR5-SNAP construct cause the lack of binding of the native chemokines, since the construct shows wild-type functionality in cell-based assays. On possibility is that the native chemokines require G-protein pre-coupling to bind with high affinity to CCR5.(11, 31) Another possibility that can explain the lack of observed native chemokine binding to CCR5 in our assay is that upon detergent solubilization, the receptor adopts a conformation that is specific only to the RANTES analogs. To discern between these different possibilities, future experiments will probe the role of G-protein pre-coupling on the ligand binding interactions of the RANTES analogs and native chemokines to CCR5-SNAP in a more native lipid environment, such as nanoscale apolipoprotein bound bilayers (NABBs),(24, 32) since CCR5 solubilized in detergent micelles cannot efficiently bind G-protein.

We generated homology models of 5P12 and 6P4 binding to CCR5 based on the crystal structure of the 5P7-CCR5 complex to elucidate the structural determinants of their different affinities.(13) Note that 5P7 and 5P12 differ by only one amino acid.(9) Fig. 6 shows the lowest energy conformations for 5P12-RANTES and 6P4-RANTES bound to CCR5. 5P12-RANTES and 6P4-RANTES adopt a conformation that closely mimics the conformation observed for 5P7 in the crystal structure. In our model, we observe that the 6P4-CCR5 complex has a negatively charged residue, Asp5 in 6P4, in the binding pocket not stabilized by a counter ion. We speculate that the lack of stabilizing counterion interactions with this residue may help to explain the lower affinity observed for 6P4 relative to 5P12.

**Figure 6.**
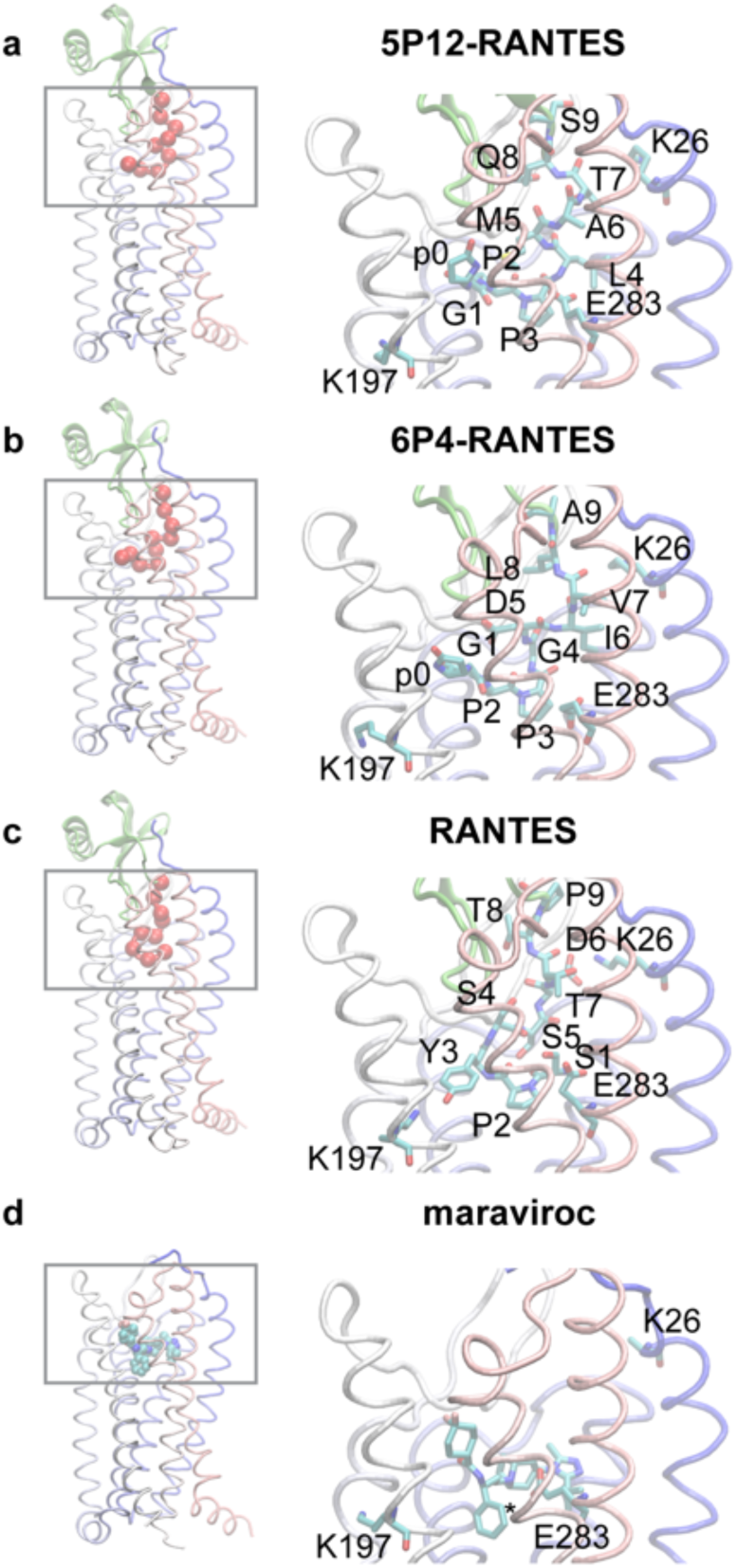
CCR5-ligand homology structural models based on the crystal structure of the CCR5-5P7 complex. Homology structural models of CCR5 bound to 5P12-RANTES (a), 6P4-RANTES (b), RANTES (c) and maraviroc (d). The structural models are based on the crystal structures of the CCR5-5P7 complex and the CCR5-maraviroc complex. The N-terminal tail residues of the chemokines are shown as red beads on a string. The remaining part of the chemokines is shown in ribbon representation colored in green. Maraviroc is shown using a blue space-filling model. CCR5 is represented as a ribbon structure with each transmembrane segment colored differently. The figure illustrates the different orientations observed for the bound chemokine analogs relative to Glu283 in CCR5. The panels on the right show the region marked within the grey rectangle in higher magnification. Chemokine residues 1-9 are labeled as well as residues K26, K197, and E283 in CCR5 using single-letter amino acid abbreviations. The pyroglutamate residue 0 of the analogs is labeled as p0.

Glu283 in CCR5 is another charged residue with no stabilizing counterions in the 5P12- and 6P4-CCR5 complex models. Zheng et al. observed that Glu283 makes a hydrogen bond with Leu4 in the 5P7-CCR5 crystal structure (13). We observe a similar hydrogen bond interaction with Glu283 and Leu4 in the 5P12-CCR5 complex model, whereas the backbone of 6P4 was removed from Glu283. Choi et al. demonstrated that Glu283 is a key residue for the anti-viral activities of 5P12 and 6P4.(33) Using the CCR5 E283A mutant, Choi et al. measured a 100-fold loss in anti-viral potency for 5P12, while for 6P4 a 100-fold increase in potency supporting the hypothesis that Glu283 destabilizes 6P4 binding to CCR5. Therefore, we hypothesize that the lower affinity observed for 6P4 in our FCCS measurements relative to 5P12 may also be due to the lack of stabilizing interactions with Glu283 in CCR5.

We also modeled RANTES binding to CCR5 (Fig. 6c) and we observed that the RANTES N-terminal tail adopts a different conformation in the ligand-binding pocket relative to that of 6P4 and 5P12. The free amino terminus of Ser1 in RANTES makes a salt bridge with Glu283 in CCR5 and we attribute the different conformations observed between RANTES and the chemokine analogs primarily to this salt bridge. Previous experimental and theoretical studies have shown that Glu283 is necessary for native chemokine binding to CCR5.(34) Moreover, the maraviroc-CCR5 crystal structure (Fig. 6d) shows that the positively-charged tropane nitrogen forms a salt bridge with Glu283 (35). Likewise, previous alanine scanning mutagenesis in CCR5 and modeling studies have shown that Glu283 is necessary for the binding of positively charged small molecule antagonists to CCR5.(36-38)

Another difference between RANTES and 5P12/6P4 in our binding models is that the hydroxyl of Tyr3 in RANTES is close to the ε-amine of Lys197 alluding to the possibility that Tyr3 forms a salt bridge with Lys197 as a tyrosinate. Lys197 in CCR5 is buried in the binding pocket with no obvious stabilizing charges in proximity. Tyr3 is close to the hydrophobic pocket that is occupied by the 1,1-difluorocyclohexane moiety in the maraviroc-CCR5 complex and the pyroglutamate residue in the 5P12 and 6P4 complexes. In contrast to the isolated Asp5 in the CCR5-6P4 complex, Asp6 in RANTES makes a salt bridge with Lys26 of CCR5.

In conclusion, we have presented methods to prepare functional, monomeric, stoichiometrically fluorophore-conjugated receptors and fluorescent chemokine ligands that provide a fundament for single-molecule studies of ligand–GPCR interactions. We demonstrate convenient quantification of nanomolar receptor concentration by FCS, and we developed FCCS saturation and competition binding assays for ligand-receptor interactions with nanomolar affinities. In a chemically defined, purified system, which is not affected by cellular heterogeneity, we have demonstrated high-affinity binding of high potency anti-HIV chemokines to naked receptor consistent with a model proposed for G-protein-biased signaling in this system.

Supporting Information. Supporting information contains supplementary text, tables, figures and references as listed in the main text. This material is available free of charge via the Internet at http://biorxiv.org.

## AUTHOR CONTRIBUTIONS

C.A.R., A.F., O.H., T.P.S., and T.H. designed the research; C.A.R., Y.A.B., M.H., E.L., H.T., M.A.K., H.F., and T.H. performed the research; J.C.P. provided new reagents; C.A.R., Y.A.B., E.L., A.F., O.H., T.P.S., and T.H. analyzed the data; and C.A.R., Y.A.B., E.L., A.F., T.P.S., and T.H. wrote the article.

## ACKNOWLEDGEMENTS

We thank the Rockefeller University Bio-Imaging Resource Center for training, assistance and discussions related to fluorescent imaging and FCCS experiments. We thank the High-Throughput and Spectroscopy Resource Center for their training and access to the LiCor instrument for NIR-immunoblots. We thank Dr. W Vallen Graham for technical assistance with the receptor purifications. E.L. was supported by the David Rockefeller Graduate program. C.A.R and M.H. were supported by the Tri-institutional Training Program in Chemical Biology T32 GM115327. C.A.R. was supported by the National Science Foundation Graduate Research Fellowship Grant DGE-1325261.

