## Supplementary Information for "High-Affinity Binding of Chemokine Analogs that Display Ligand Bias at the HIV-1 Co-receptor CCR5"

### SUPPLEMENTARY TEXT

*Discussion of variations in correlation function amplitude, diffusion time and molecular brightness*

We observed well-to-well variations in the correlation function amplitudes for samples with identical concentrations that are show in parallel variations in the diffusion times and molecular brightness (Supplementary Figure S10-S9). Variations in the diffusion time can either be attributed to changes in the diffusion coefficient or the confocal volume. Since all samples were prepared using the same buffer composition(1), we can rule out changes in the diffusion coefficient. Thus, we attributed the non-random variations in $A$ to variations in the confocal volume when the underlying diffusion coefficient is constant. The confocal volume most likely changes as a result of optical aberrations due to uneven glass substrates in the microtiter plates we used. With increasing confocal volume, the diffusion time and number of particles should increase, whereas the molecular brightness and the amplitudes should decrease. We also observed similar non-random variations in molecular brightness (Supplementary Fig. S10), triplet state transitions (Supplementary Fig. S11) and blinking transitions (Supplementary Fig. S12) of the three components. The blinking times should not vary with confocal volume changes, and it is likely that the blinking term actually fits a secondary diffusion component of the ligand. The impact of the blinking term variation on the overall calculations is small, and we decided not to investigate this effect further. However, we reduced the impact of these confocal volume variations on the calculations of the ligand binding equilibrium by using averaged receptor concentrations for each individual experimental data set and by calculating the fractional occupancy from $A_X/A_L$, which cancels out correlated variations of the confocal volumes.

*Native chemokines and sCD4/gp120 complex do not compete with 5P12 and 6P4*

We also performed saturation binding measurements with RANTES-647 and CCR5-SNAP-488 but we did not observe significant binding to the receptor (Supplementary Fig. 13). We hypothesized that the affinity of RANTES-647 is too low for saturation binding detection by FCCS, so we performed competition binding with 5P12-647 or 6P4-647 with unlabeled RANTES (Supplementary Fig. 14). To our surprise, RANTES could not displace either 5P12-647 or 6P4-647 at up to 10 µM concentration. Also MIP-1α, another native CCR5 chemokine ligand, was ineffective at displacing either labeled ligand up to 10 µM concentration (Supplementary Fig. 15). The lack of competition by the native chemokines is unexpected since we showed earlier that they activate Gαi signaling through CCR5-SNAP in live cells, but consistent with the notion that these native chemokines only bind to the G-protein pre-coupled receptor.(2) We then performed competition binding with human soluble CD4 (sCD4) complexed to 2G12-purified monomeric gp120 (sCD4/gp120) in the presence of 5P12-647 or 6P4-647. We also did not observe a reduction in 5P12-647 and 6P4-647 binding to CCR5-SNAP-488 up to 1 µM of sCD4/gp120 (Supplementary Fig. 16). Our results show that the native chemokines and sCD4/gp120 cannot displace the RANTES analogues from purified CCR5-SNAP-488.

### SUPPLEMENTARY FIGURES

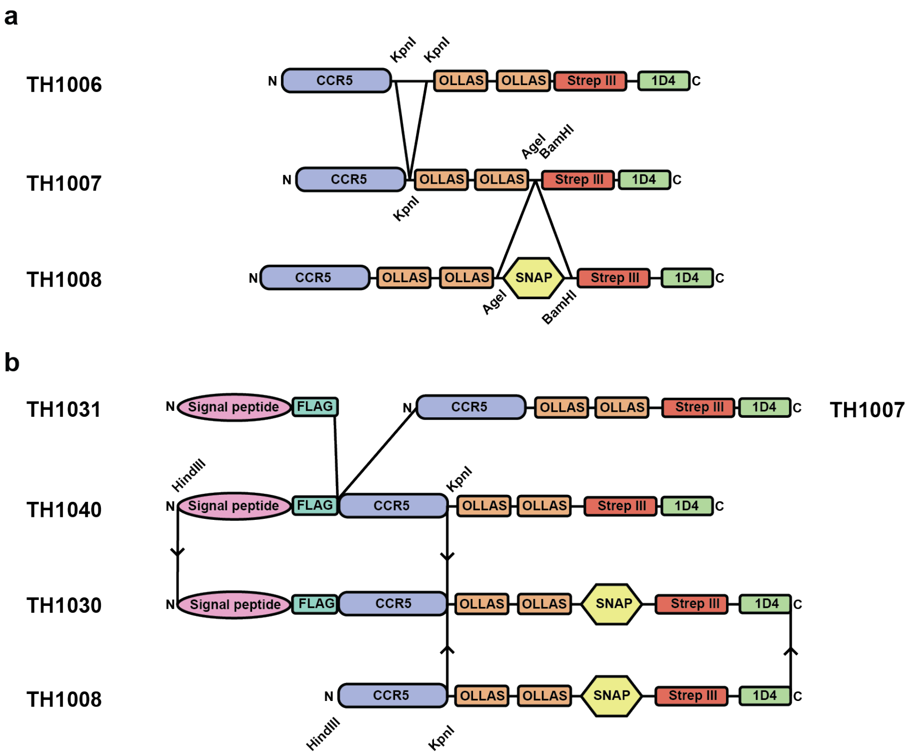

Supplementary Figure S1. Schematic showing the different CCR5 constructs employed to generate the CCR5-SNAP construct (TH1030) employed in this study. Each construct shows the functional tags encoded and relevant restriction sites that were employed to generate new constructs. a) shows the synthesis of TH1008 from TH1006 as a starting point. b) shows the synthesis of TH1030 from TH1008 and TH1040 which was designed from TH1031 and TH1007. Lines indicate additions or deletions within constructs and adjacent to those the restriction enzymes employed for each step.

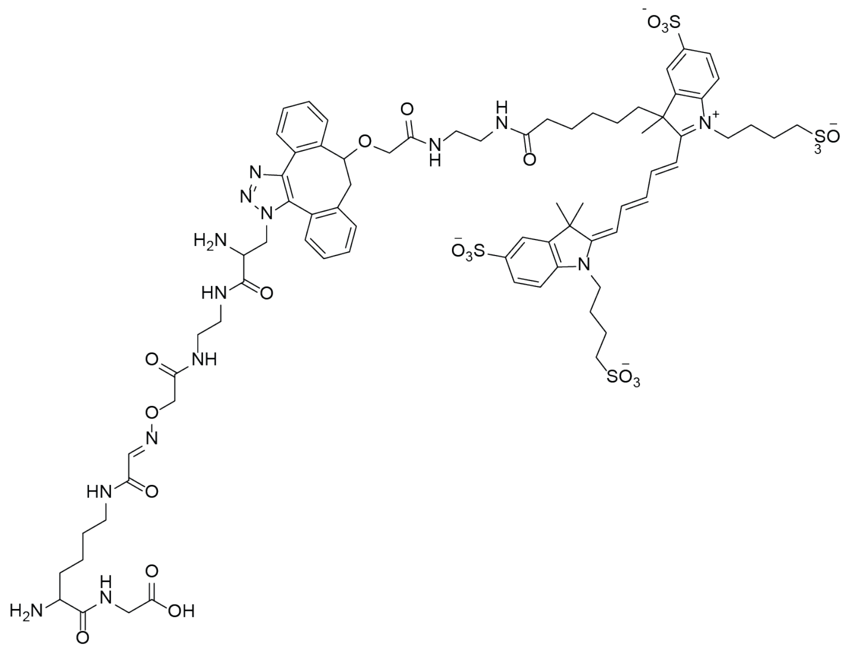

Supplementary Figure S2. Synthesis of fluorescently-labeled chemokines. The chemokines (residues 1–68) were synthesized by native chemical ligation of two similar sized fragments. The C-terminal fragment carried an additional K(Alexa647)G sequence with the structure shown here.

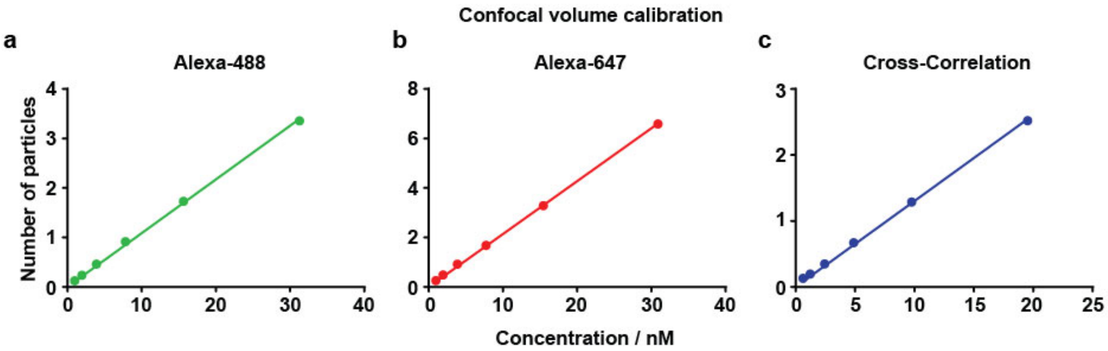

Supplementary Figure S3. Confocal volume determination. Number of particles as a function of concentration of Alexa-488 (a), Alexa-647 (b), or Alexa-488/Alexa-647 dual-labeled 40-base pair oligonucleotide (c). Data points were fitted using a linear equation where the $y$-intercept is set to 0. Confocal volume size is derived by dividing the slope by Avogadro's number. We calculated a confocal volume of 0.18 fL for 488-nm excitation, 0.35 fL for 633 nm excitation, and 0.22 fL for the overlap between the two laser excitations.

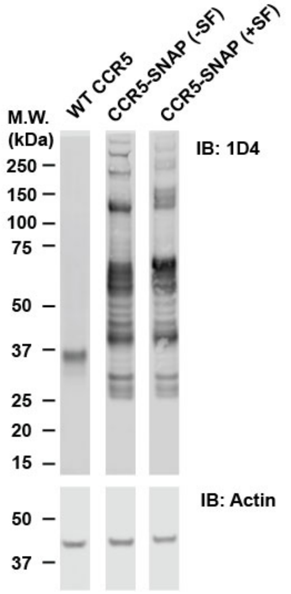

Supplementary Figure S4. Qualitative expression comparison between wild-type CCR5 and CCR5-SNAP constructs. SDS-PAGE western blots of wild-type CCR5, CCR5-SNAP lacking the N-terminal signal peptide and FLAG epitope [CCR5-SNAP (-SF)] and CCR5-SNAP encoding the N-terminal signal peptide and FLAG epitope [CCR5-SNAP (+SF)]. 1,000,000 cells per construct were transfected with 20 ng of plasmid and then solubilized in 1% DDM (w/v) 24 hours post-transfection. CCR5 was detected using an anti-1D4 antibody. Actin was detected as a loading control. Wild-type CCR5 runs as a single band at approximately 35 kDa and both full-length CCR5-SNAP (-SF) and CCR5-SNAP (+SF) run at an approximate molecular weight of 70 kDa. Both CCR5-SNAP constructs express at higher levels than wild-type CCR5 but also show the presence of several N-terminal receptor truncations.

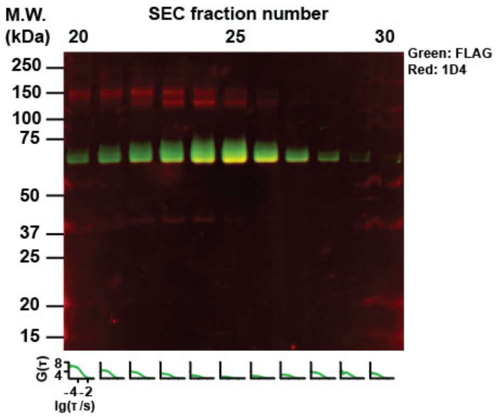

Supplementary Figure S5. Characterization of CCR5-SNAP-488 SEC fractions 20 to 30. SDS-PAGE and NIR western blot (WB) of SEC fractions 20 to 30. Full-length CCR5-SNAP (~70 kDa, yellow band) was detected using antibodies against the 1D4 (red) and FLAG (green) epitopes. Lower inset shows the FCS auto-correlation curves obtained from the same SEC fractions tested by WB.

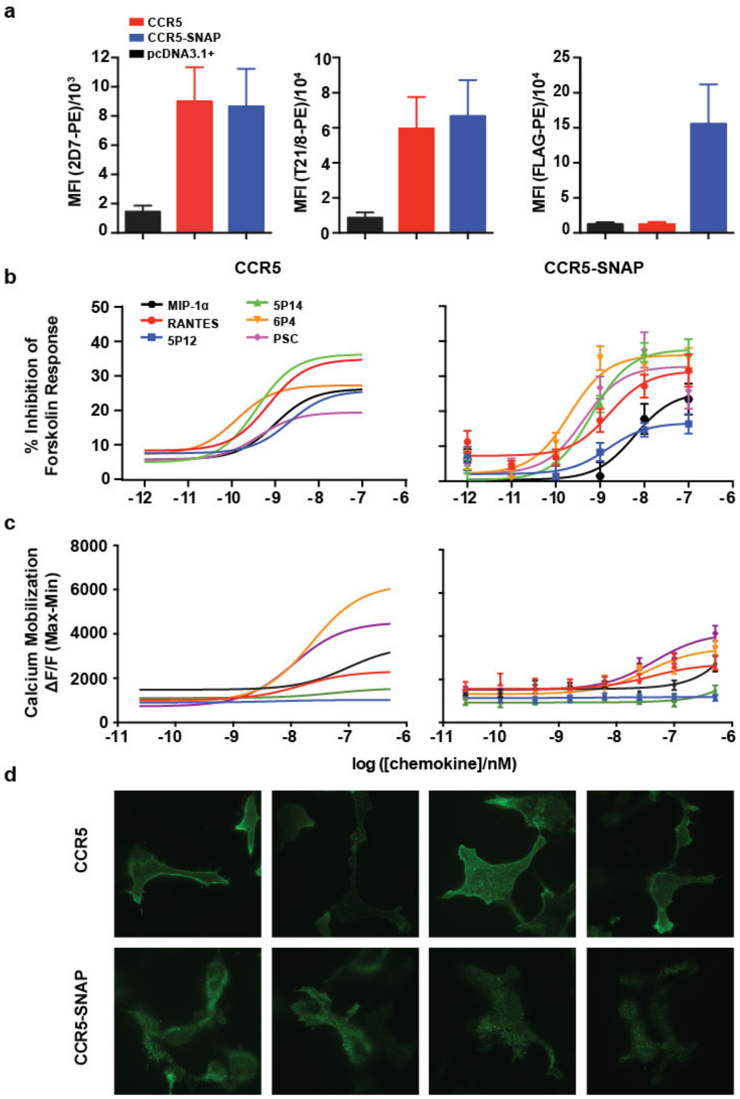

Supplementary Figure S6. Functional characterization of CCR5-SNAP (+SF). (a) Cell surface quantification by flow cytometry of wild-type CCR5 and CCR5-SNAP (+SF), simply referred to as CCR5-SNAP, in HEK293T cells stained with anti-CCR5 (clone 2D7), anti-CCR5 (clone T21/8), or anti-FLAG coupled to phycoerythrin (PE). Data are mean of four independent experiments ± S.E.M. (b) Percent inhibition of forskolin-induced cAMP production by wild-type CCR5 and CCR5-SNAP in response to increasing concentrations of MIP-1α (black), RANTES (red), 5P12 (blue), 5P14 (green), 6P4 (yellow), and PSC (violet). Fits are only shown for wild-type CCR5 as a direct comparison (Lorenzen et al. 2017). Data points are mean from 3 independent experiments performed in triplicate ± S.E.M. (c) Calcium flux by wild-type CCR5 and CCR5-SNAP in response to increasing concentrations of the indicated native and chemokine analogues. Data points represent the mean of the maximum fluorescence minus basal fluorescence from three independent experiments performed in triplicate ± S.E.M. (d) Fluorescent images of HEK293T cells expressing wild-type CCR5 or CCR5-SNAP imaged using total internal reflection microscopy. CCR5 was detected using the anti-1D4 mouse antibody and Alexa-488 conjugated anti-mouse secondary antibody.

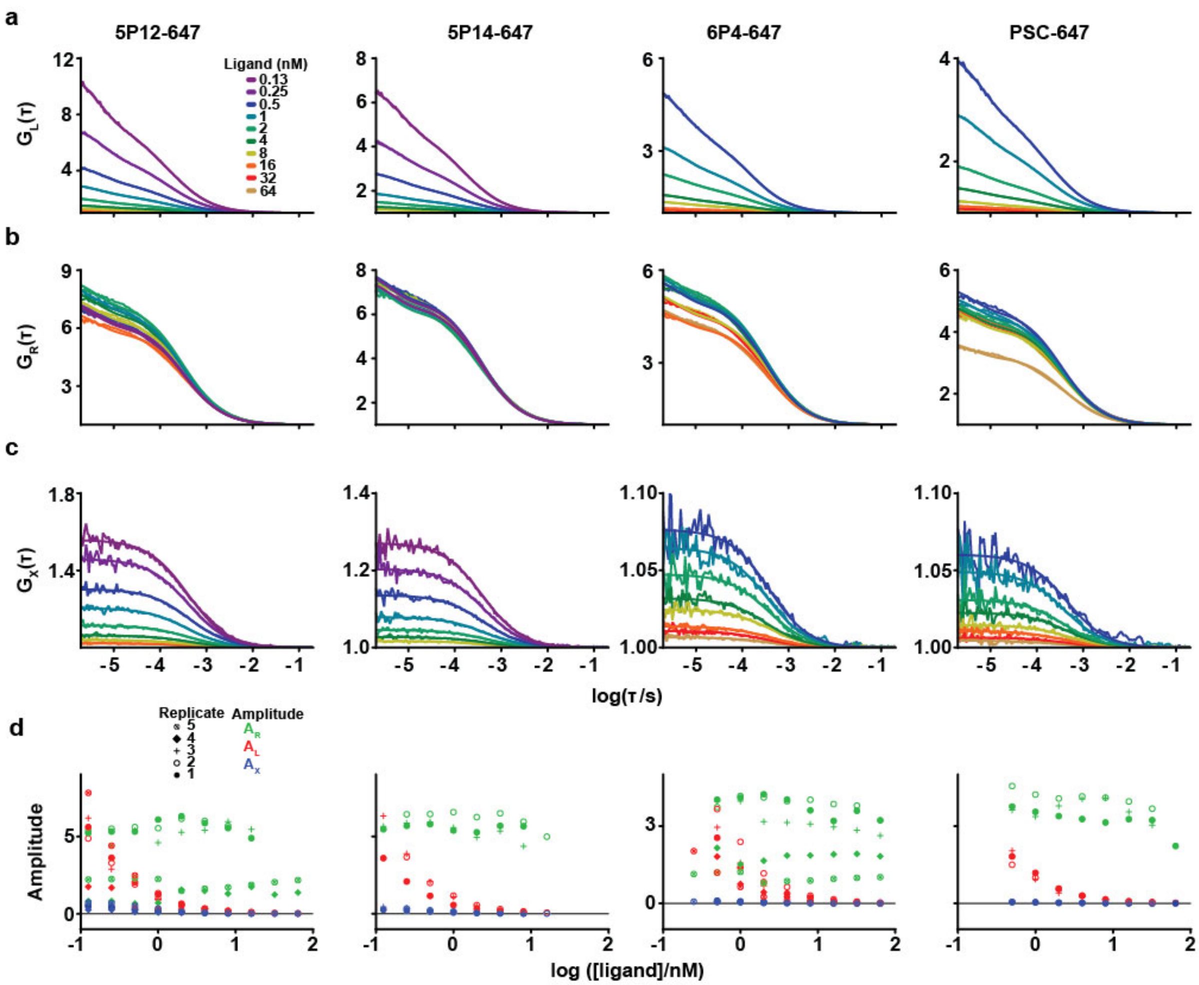

Supplementary Figure S7. Correlation traces from saturation binding of Alexa-647 labeled RANTES analogues to CCR5-SNAP-488 by FCCS. (a) Autocorrelation traces and fits for 5P12-647, 5P14-647, 6P4-647, and PSC-647 binding to CCR5-SNAP-488 at various ligand concentrations. Ligand autocorrelation traces were fitted to a one-component with 3D translational diffusion with independent triplet state and blinking transitions. (b) CCR5-SNAP-488 autocorrelation traces and fits for different concentrations of 5P12-647, 5P14-647, 6P4-647, and PSC-647. CCR5-SNAP-488 correlation traces were fitted to a one-component with 3D translational diffusion with triplet state transitions. (c) Receptor-ligand complex cross-correlation traces and fits for different concentrations of 5P12-647, 5P14-647, 6P4-647, and PSC-647. Cross-correlation traces were fitted to a one-component with 3D translational diffusion. Autocorrelation and cross-correlation amplitudes and traces shown in (a-c) are representative from one independent experiment for each Alexa-647 labeled RANTES analogue. Correlation amplitudes for CCR5-SNAP-488 ($A_R$, green), Alexa-647 labeled analogue ($A_L$, red), and ligand-receptor complex ($A_X$,blue) at various ligand concentrations from at least 3 independent experiments. The black line at $A = 0$ is shown to clearly illustrate the changes that occur in correlation amplitudes across concentrations.

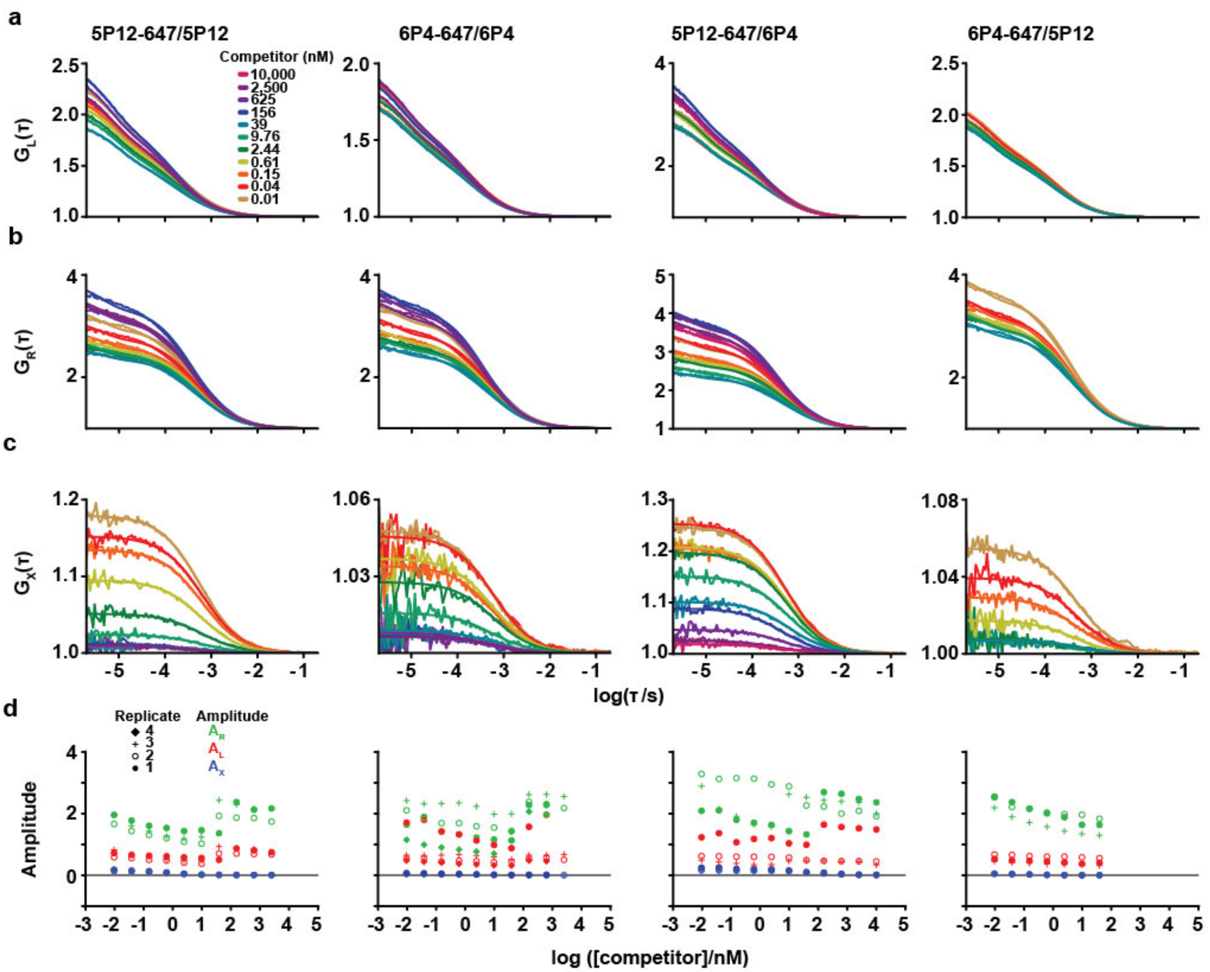

Supplementary Figure S8. Correlation traces from competition binding of Alexa-647 labeled RANTES analogues with non-labeled analogues by FCCS. (a) Autocorrelation traces and fits for 5P12-647 in competition with 5P12 or 6P4 and 6P4-647 in competition with 6P4 or 5P12 at various competitor concentrations. Ligand autocorrelation traces were fitted to a one-component with 3D translational diffusion with independent triplet and blinking state transitions. (b) CCR5-SNAP-488 autocorrelation traces and fits for 5P12-647 in competition with 5P12 or 6P4 and 6P4-647 in competition with 6P4 or 5P12. CCR5-SNAP-488 correlation traces were fitted to a one-component with 3D translational diffusion with triplet state transitions. (c) Receptor-ligand complex cross-correlation traces and fits for 5P12-647 in competition with 5P12 or 6P4 and 6P4-647 in competition with 6P4 or 5P12. Cross-correlation traces were fitted to a one-component with 3D translational diffusion. Autocorrelation and cross-correlation amplitudes and traces shown in (a-c) are representative from one independent experiment. Correlation amplitudes for CCR5-SNAP-488 ($A_R$, green), Alexa-647 labeled analogue ($A_L$, red), and ligand-receptor complex ($A_X$,blue) at various competitor concentrations from at least 3 independent experiments. The black line at $A = 0$ is shown to clearly illustrate the changes that occur in correlation amplitudes across competitor concentrations.

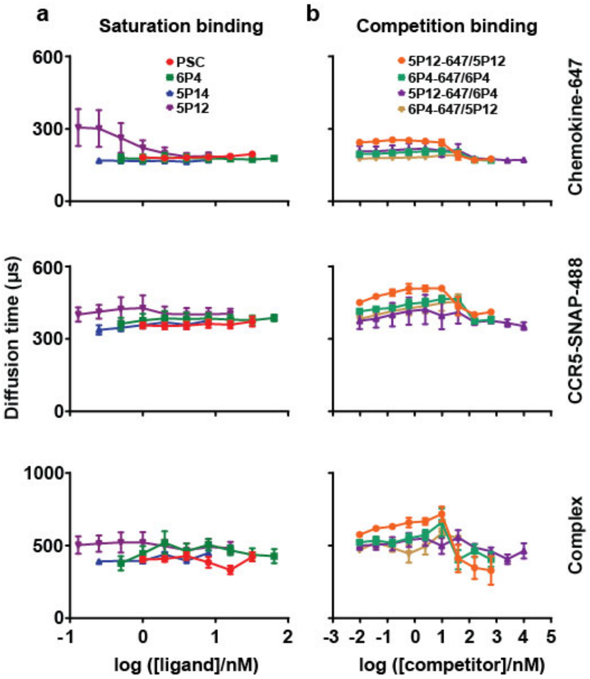

Supplementary Figure S9. Diffusion time dependency on ligand or competitor concentration. Average diffusion times ($\tau_D$) in μs determined from fitting autocorrelation and cross-correlation curves for the Alexa-647 labeled RANTES analogues, CCR5-SNAP-488, and the receptor-ligand complex as a function of labeled chemokine or competitor concentration. Values are averages from at least 3 independent saturation binding (a) or competition binding (b) experiments and errors are the S.E.M.

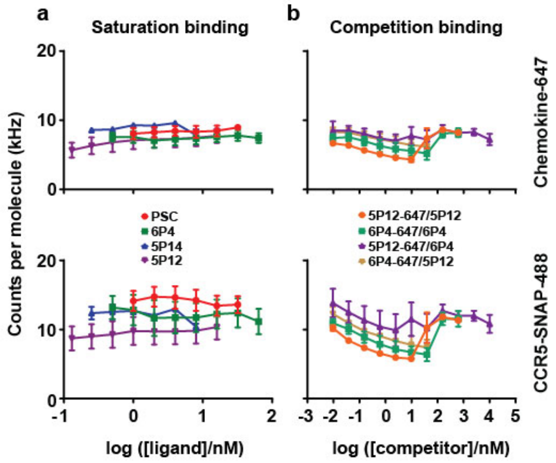

Supplementary Figure S10. Brightness dependency on ligand or competitor concentration. Average brightness values measured as counts per molecule (kHz) for the RANTES analogues labeled with Alexa-647 and CCR5-SNAP-488 as a function of labeled chemokine or competitor concentration. Values are averages from at least 3 independent saturation binding (**a**) or competition binding (**b**) experiments and errors are the S.E.M.

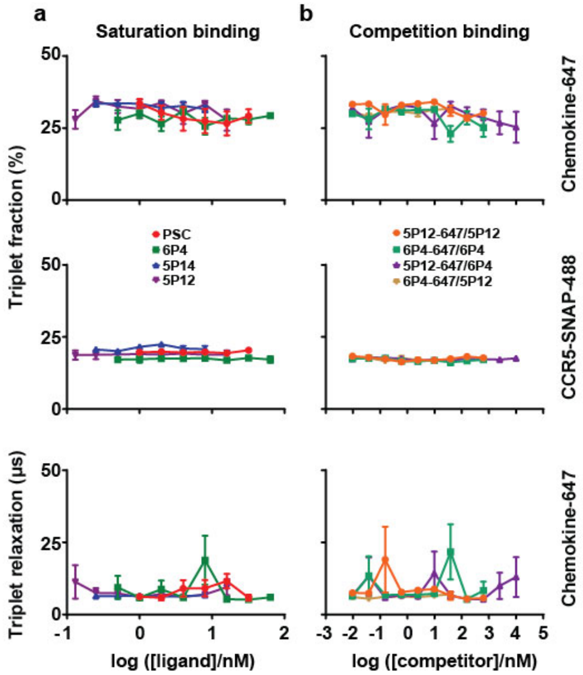

Supplementary Figure S11. Triplet state dependency on ligand or competitor concentration. Average triplet state fraction and relaxation times in μs determined from fitting autocorrelation and cross-correlation curves for the Alexa-647 labeled RANTES analogues and CCR5-SNAP-488 as a function of labeled chemokine or competitor concentration. CCR5-SNAP-488 triplet relaxation time was fixed to 4 μs in all experiments. Values are averages from at least 3 independent saturation binding (**a**) or competition binding (**b**) experiments and errors are the S.E.M.

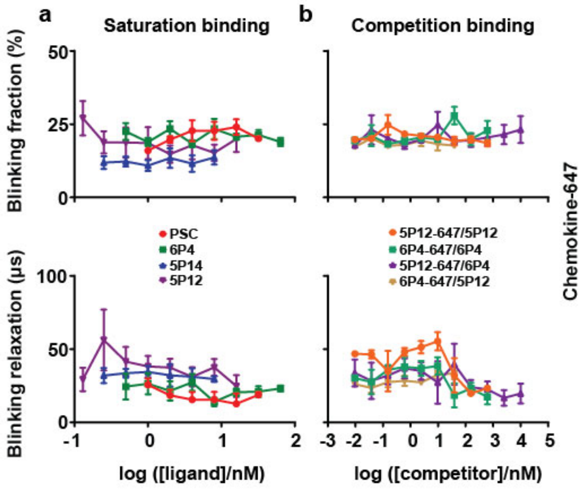

Supplementary Figure S12. Blinking state dependency on ligand or competitor concentration. Average blinking state fraction and relaxation time in μs determined from fitting autocorrelation curves for the Alexa-647 labeled RANTES analogues as a function of labeled chemokine or competitor concentration. Values are averages from at least 3 independent saturation binding (a) or competition binding (b) experiments and errors are the S.E.M.

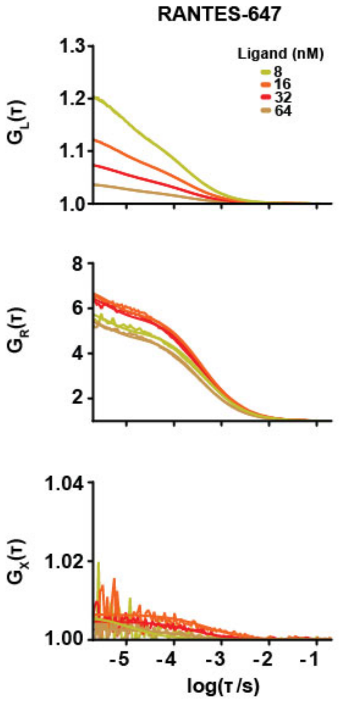

Supplementary Figure S13. Saturation binding of RANTES-647 with CCR5-SNAP-488. RANTES-647 and CCR5-SNAP-488 auto-correlation traces and receptor-ligand complex cross-correlation traces at various concentrations of RANTES-647. Ligand correlation traces were fitted with one-component with 3D translational diffusion with independent triplet and blinking state transitions. CCR5-SNAP-488 correlation traces were fitted to a one-component with 3D translational diffusion with triplet state transitions. Cross-correlation traces were fitted to a one-component with 3D translational diffusion.

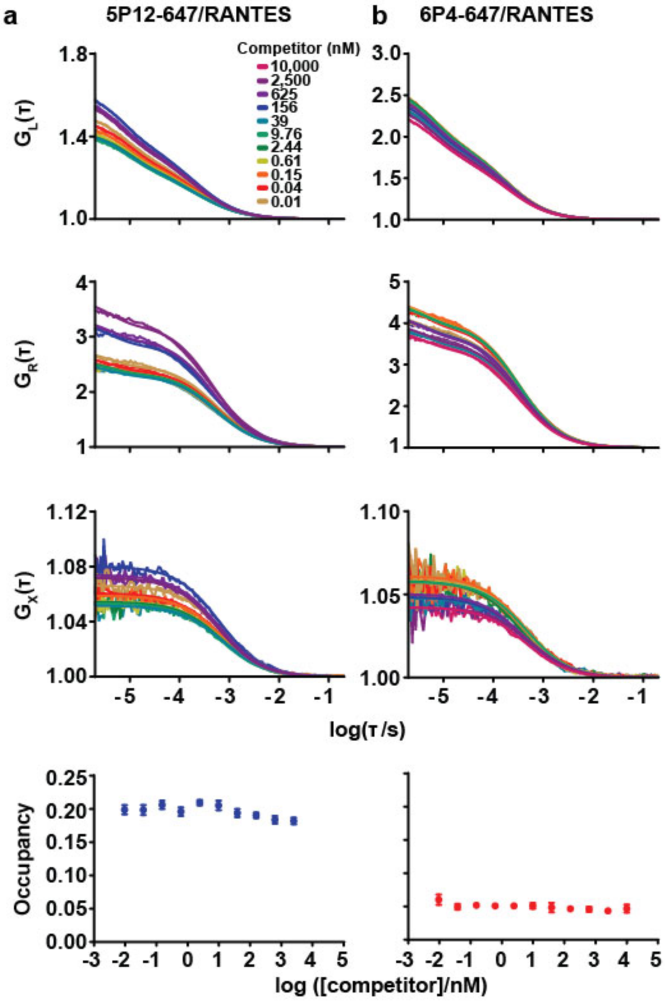

Supplementary Figure S14. Competition binding of 5P12-647 and 6P4-647 with RANTES. (a) 5P12-647 and 6P4-647 auto-correlation traces and fits for all concentrations of RANTES tested. Ligand correlation traces were fitted with a one-component with 3D translational diffusion with independent triplet and blinking state transitions. (b) CCR5-SNAP-488 auto-correlation traces and fits in the presence of 5P12-647 and 6P4-647 at different concentrations of RANTES. CCR5-SNAP-488 correlation traces were fitted to a one-component with 3D translational diffusion with triplet state transitions. (c) Receptor-ligand complex cross-correlation traces and fits for 5P12-647 and 6P4-647 at different concentrations of RANTES. Cross-correlation traces were fitted to a one-component with 3D translational diffusion. (d) Competition binding isotherms for 5P12-647 and 6P4-647 at various concentrations of RANTES. Data points represent the mean from at least 30 individual FCCS measurements and their associated errors.

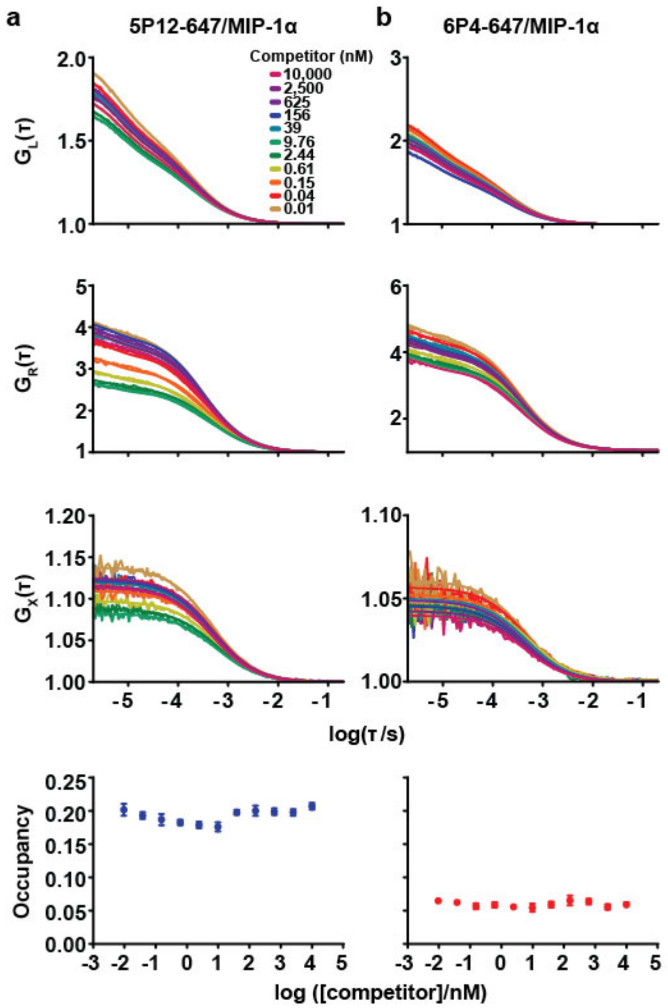

Supplementary Figure S15. Competition binding of 5P12-647 and 6P4-647 with MIP-1α. (a) 5P12-647 and 6P4-647 auto-correlation traces and fits for all concentrations of MIP-1α tested. Ligand correlation traces were fitted to a one-component with 3D translational diffusion with independent triplet and blinking state transitions. (b) CCR5-SNAP-488 auto-correlation traces and fits in the presence of 5P12-647 and 6P4-647 at different concentrations of MIP-1α. CCR5-SNAP-488 correlation traces were fitted to a one-component with 3D translational diffusion with triplet state transitions. (c) Receptor-ligand complex cross-correlation traces and fits for 5P12-647 and 6P4-647 at different concentrations of MIP-1α. Cross-correlation traces were fitted to a one-component with 3D translational diffusion. (d) Competition binding isotherms for 5P12-647 and 6P4-647 at various concentrations of MIP-1α. Data points represent the mean from at least 30 individual FCCS measurements and their associated errors.

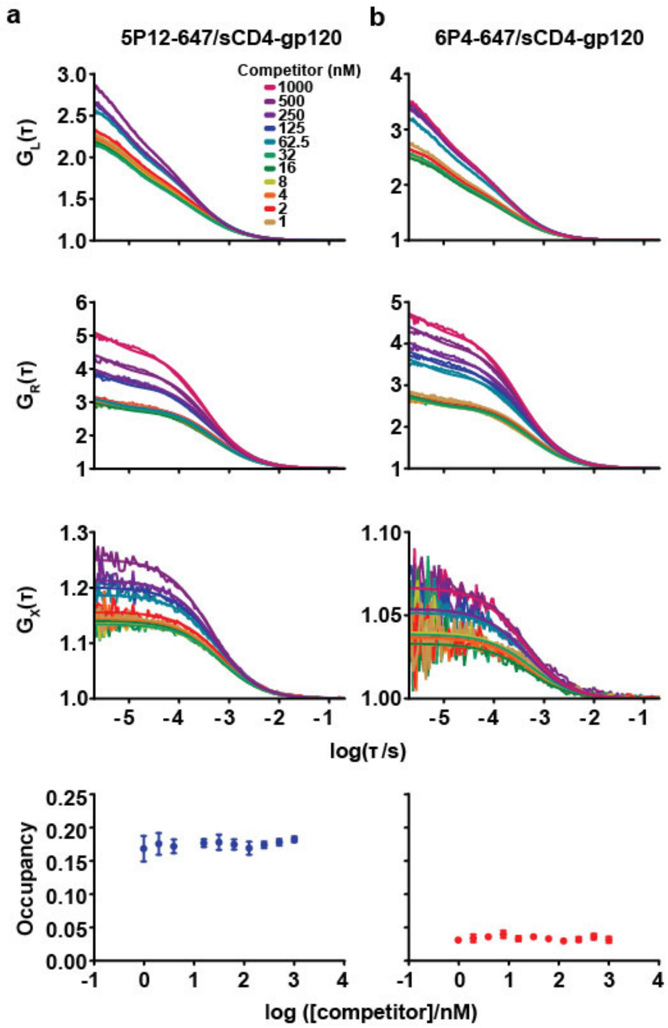

Supplementary Figure S16. Correlation traces for 5P12-647 and 6P4-647 in competition with the sCD4-gp120 complex. (a) 5P12-647 and 6P4-647 auto-correlation traces and fits for all concentrations of sCD4-gp120 complex tested. Ligand correlation traces were fitted to a one-component with 3D translational diffusion with independent triplet and blinking state transitions. (b) CCR5-SNAP-488 auto-correlation traces and fits in the presence of 5P12-647 and 6P4-647 at different concentrations of sCD4-gp120 complex. CCR5-SNAP-488 correlation traces were fitted to a one-component with 3D translational diffusion with triplet state transitions. (c) Receptor-ligand complex cross-correlation traces and fits for 5P12-647 and 6P4-647 at different concentrations of sCD4-gp120 complex. Cross-correlation traces were fitted to a one-component with 3D translational diffusion. (d) Competition binding isotherms for 5P12-647 and 6P4-647 at various concentrations of sCD4-gp120 complex. Data points represent the mean from at least 30 individual FCCS measurements and their associated errors.

### SUPPLEMENTARY TABLES

| Ligand | N-terminal Sequence 0-1-2-3-4-5-6-7-8-9 | Anti-HIV potency (pM) |
|---|---|---|
| RANTES | S-P-Y-S-S-D-T-T-P | ~ 1000000 |
| PSC-RANTES | β-γ-δ-S-S-D-T-T-P | 25 |
| 6P4-RANTES | α-G-P-P-G-D-I-V-L-A | 21 |
| 5P12-RANTES | α-G-P-P-L-M-A-T-Q-S | 28 |
| 5P14-RANTES | α-G-P-P-L-M-S-L-Q-V | 26 |
| MIP-1α | S-L-A-A-D-T-P-T-A | ~ 1000000 |

Supplementary Table S1. Peptide sequences and pharmacological properties of RANTES and some of its peptide analogues. The RANTES analogues investigated in this report have several amino acid mutations within the first 9 amino acids of the native N-terminus. The RANTES and MIP-1α N-terminus sequences are shown for comparison. PSC-RANTES is a chemically modified analogue where PSC is (*N*-nonanoyl)-*des*-Ser-[L-thioproline, L-2-cyclohexylglycine]. The phage-display derived analogs have 10 amino acids at the N-terminus. The symbol α corresponds to pyroglutamate, β corresponds to *N*-nonanoyl, γ corresponds to L-thioproline, and δ corresponds to L-2-cyclohexylglycine. RANTES has very low anti-HIV potency while the RANTES analogues display picomolar inhibition potencies.(3)

| | TH1006 | |
|---|---|---|
| | ( A S S R N N E F K L P A G D D T ) | Untranslated |
| | gctagctctagaaataatgaattcaagcttcctgcaggagatgatacc | DNA |
| 5' Adapter | NheI-XbaI-EcoRI-HindIII-SbfI-Kozak | Endonuclease |
| hCCR5 | (1) MDYQVS … EISVGL (352) | Amino acid |
| Linker | TETAST | Amino acid |
| | G T | Amino acid |
| KpnI | ggtacc | DNA |
| Ollas (2-14) | GFANELGPRLMGK | Amino acid |
| Ollas | SGFANELGPRLMGK | Amino acid |
| Spacer4 | SGGG | Amino acid |
| | D I | Amino acid |
| EcoRV | gatatc | DNA |
| | T G A G S | Amino acid |
| AgeI-BamHI | accggtgctggatcc | DNA |
| | G S P | Amino acid |
| BstEII | gggtcacct | DNA |
| Strep-III | WSHPQFEKGGGSGGGSGGGSWSHPQFEK | Amino acid |
| 1D4 | DEASTTVSKTETSQVAPA* | Amino acid |
| | ctcgaggcggccgc | DNA |
| 3'adapter | XhoI-NotI | Endonuclease |

Supplementary Table S2. Functional groups encoded in TH1006. Amino acid and/or DNA sequences are shown for each functional group. In the case of DNA sequences, restriction sites are color-coded where relevant. Numbers in parentheses indicate the number position of the amino acid. The * symbol represents the stop codon in the amino acid sequence.

| TH1031 | | |
|---|---|---|
| 5' Adapter | tctagagctagcgagtcttcaagcttcctgcaggagatgataccgccgccacc<br>XbaI-NheI-HindIII-MlyI-SbfI-Kozak | DNA<br>Endonuclease |
| Signal Peptide | MRLCIPQVLLALFLSMLTGPGEG | Amino acid |
| FLAG | DYKDDDDK | Amino acid |
| CCR5 (1-7) | (1) MDYQVSS (7) | Amino acid |
| MlyI | G G T Q<br>ggcgggactcaa | Amino acid<br>DNA |
| AgeI-BamHI | T G A G S<br>accggtgctggatcc | Amino acid<br>DNA |
| KpnI | G T<br>ggtacc | Amino acid<br>DNA |
| Ollas | SGFANELGPRLMGK | Amino acid |
| Spacer4 | SGGG | Amino acid |
| EcoRV | D I<br>gatatc | Amino acid<br>DNA |
| Spacer20 | SGGGSGGGSGGGSGGGSGGG | Amino acid |
| Ollas | SGFANELGPRLMGK | Amino acid |
| 1D4 | DEASTTVSKTETSQVAPA* | Amino acid |
| 3' Adapter | ctcgaggcggccgctcccggg<br>XhoI-NotI-XmaI | DNA<br>Endonuclease |

Supplementary Table S3. Functional groups encoded in TH1031. Amino acid and/or DNA sequences are shown for each functional group. In the case of DNA sequences, restriction sites are color-coded where relevant. Numbers in parentheses indicate the number position of the amino acid. The * symbol represents the stop codon in the amino acid sequence.

| | Primer Nucleotide Sequence |
|---|---|
| Forward | 5'-GCTTCCTGCAGGAGATGATACC-3’ |
| Reverse | 5'-GCTGGACACCTGGTAATCCAT-3' |

Supplementary Table S4. Primer nucleotide sequences. Primers were employed to amplify the signal peptide and FLAG sequence from TH1031.

| | ***cAMP inhibition*** | | | | |
|---|---|---|---|---|---|
| **Receptor** | **Ligand** | **N** | **$EC_{50}$ (nM)** | **$pEC_{50}$ ± S.E.M.** | **[illegible]$E_{max}$ ± S.E.M.** |
| CCR5 | RANTES | 3 | 0.8 | 9.2 ± 0.2 | 26 ± 2 |
| | MIP-1✓ | 3 | 1.3 | 9.3 ± 0.5 | 21 ± 4 |
| | 5P12-RANTES | 3 | 2.4 | 8.7 ± 0.1 | 17 ± 6 |
| | 5P14-RANTES | 3 | 0.5 | 9.4 ± 0.2 | 32 ± 3 |
| | 6P4-RANTES | 3 | 0.1 | 9.9 ± 0.1 | 20 ± 1 |
| | PSC-RANTES | 3 | 0.7 | 9.2 ± 0.1 | 17 ± 3 |
| CCR5-SNAP | RANTES | 3 | 1.7 | 9.8 ± 0.1 | 24 ± 5 |
| | MIP-1✓ | 3 | 8.8 | 8.2 ± 0.2 | 24 ± 7 |
| | 5P12-RANTES | 3 | 1.9 | 8.9 ± 0.3 | 15 ± 3 |
| | 5P14-RANTES | 3 | 0.9 | 9.2 ± 0.2 | 38 ± 6 |
| | 6P4-RANTES | 3 | 0.2 | 9.7 ± 0.1 | 34 ± 1 |
| | PSC-RANTES | 3 | 0.4 | 9.4 ± 0.1 | 31 ± 7 |

Supplementary Table S5. CCR5 and CCR5-SNAP cAMP inhibition parameters. $EC_{50}$, $pEC_{50}$, and $\Delta E_{max}$ values derived from fitting the dose-response curves for CCR5 and CCR5-SNAP. Dose-response curves were fitted in Prism with a three-parameter logistic equation. Values are the average from three independent experiments and associated errors are ± S.E.M.

| | ***Calcium flux*** | | | | |
|---|---|---|---|---|---|
| **Receptor** | **Ligand** | **N** | **$EC_{50}$ (nM)** | **$pEC_{50}$ ± S.E.M.** | **$\Delta E_{max}$ ± S.E.M.** |
| CCR5 | RANTES | 3 | 14 | 7.8 ± 0.2 | 13000 ± 1600 |
| | MIP-1✓ | 3 | 96 | 7.0 ± 0.3 | 20000 ± 3900 |
| | 5P12-RANTES | 3 | ND | ND | ND |
| | 5P14-RANTES | 3 | ND | ND | ND |
| | 6P4-RANTES | 3 | 23 | 7.6 ± 0.1 | 53000 ± 2600 |
| | PSC-RANTES | 3 | 11 | 7.9 ± 0.1 | 38000 ± 2200 |
| CCR5-SNAP | RANTES | 3 | 40 | 7.3 ± 0.4 | 12000 ± 3000 |
| | MIP-1✓ | 3 | ND | ND | ND |
| | 5P12-RANTES | 3 | ND | ND | ND |
| | 5P14-RANTES | 3 | ND | ND | ND |
| | 6P4-RANTES | 3 | 34 | 7.5 ± 0.3 | 21000 ± 3400 |
| | PSC-RANTES | 3 | 45 | 7.4 ± 0.2 | 26000 ± 3300 |

Supplementary Table S6. CCR5 and CCR5-SNAP calcium flux fitted parameters. $EC_{50}$, $pEC_{50}$, and $\Delta E_{max}$ values derived from fitting the dose-response curves for CCR5 and CCR5-SNAP. Dose-response curves were fitted in Prism with a three-parameter logistic equation. Values are the average from three independent experiments and associated errors are ± S.E.M. ND (not determined) is reported when the fit did not yield a sensible result or the fit did not converge.
